# Reduced Liver Mitochondrial Energy Metabolism Impairs Food Intake Regulation Following Gastric Preloads and Fasting

**DOI:** 10.1101/2024.10.24.620086

**Authors:** Michael E. Ponte, John C. Prom, Mallory A. Newcomb, Annabelle B. Jordan, Lucas L. Comfort, Jiayin Hu, Patrycja Puchalska, Caroline E. Geisler, Matthew R. Hayes, E. Matthew Morris

## Abstract

**Objective:** The capacity of the liver to serve as a peripheral sensor in the regulation of food intake has been debated for over half a century. The anatomical position and physiological roles of the liver suggest it is a prime candidate to serve as an interoceptive sensor of peripheral tissue and systemic energy state. Importantly, maintenance of liver ATP levels and within-meal food intake inhibition is impaired in human subjects with obesity and obese pre-clinical models. Previously, we have shown decreased hepatic mitochondrial energy metabolism (i.e., oxidative metabolism & ADP-dependent respiration) in male liver-specific, heterozygous PGC1a mice results in increased short-term diet-induced weight gain with increased within meal food intake. Herein, we tested the hypothesis that decreased liver mitochondrial energy metabolism impairs meal termination following nutrient oral pre-loads.

**Methods:** Liver mitochondrial respiratory response to changes in ΔG_ATP_ and adenine nucleotide concentration following fasting were examined in male liver-specific, heterozygous PGC1a mice. Further, food intake and feeding behavior during basal conditions, following nutrient oral pre-loads, and following fasting were investigated.

**Results:** We observed male liver-specific, heterozygous PGC1a mice have reduced mitochondrial response to changes in ΔG_ATP_ and tissue ATP following fasting. These impairments in liver energy state are associated with larger and longer meals during chow feeding, impaired dose-dependent food intake inhibition in response to mixed and individual nutrient oral pre-loads, and greater acute fasting-induced food intake.

**Conclusion:** These data support previous work proposing liver-mediated food intake regulation through modulation of peripheral satiation signals.

## 1. Introduction

The increased food intake associated with weight gain occurs through an imbalance of external obesogenic food cues and interoceptive satiation signals [1–4]. Predisposition to weight gain can result from a hypersensitivity to hunger interoceptive signals and/or insensitivity to satiety interoceptive signals [5]. Gastrointestinal (GI)-derived interoceptive satiation signals in the form of mechanoreception, neuropeptides, and neurotransmitters communicate nutrient-state dependent information primarily through vagal afferent neurons to the brain to regulate feeding behavior (reviewed [6–10]). These signals relay the size and nutritive composition of the material in the GI tract in a temporospatial manner, culminating in meal termination [11–17]. Reductions in the magnitude of these satiation signals delays meal termination, resulting in larger within-meal food intake, overall greater energy intake, leading over time to subsequent weight gain [18]. Importantly, human subjects with overweight/obesity and diet-induced obese pre-clinical rodent models have decreased satiation signaling following overfeeding [19], reduced vagal afferent expression of satiation signal receptors [20], reduced secretion and/or response to GI-derived satiation signals [21–30], decreased satiation in response to gastric and intestinal nutrients [31], reduced vagal afferent excitability in response to nutrients [32], and larger gastric volume and faster gastric emptying [33–35]. Collectively, these maladaptive phenotypes contribute to the reduction of within-meal food intake inhibition. While visceral vagal afferents are sensitive to GI factors to induce meal termination, absorbed nutrients and hormones generate signals originating from hepatic portal vein and liver parenchyma to activate hepatic vagal afferents in the regulation of food intake.

Due to the liver’s numerous catabolic/anabolic metabolic processes and location in the splanchnic circulation, considerable changes in energy demand occur during transitions from prandial to postprandial states [36]. However, hepatocytes cannot store high energy phosphate as phosphor-creatine like skeletal muscle [37; 38]. Thus, during high energy demand the liver can only maintain ATP levels (ATP homeostasis) by increasing mitochondrial energy metabolism ([MEM], i.e., oxidative metabolism, ADP-dependent respiration). As such, the liver is anatomically and physiologically well placed to serve as an interoceptive sensor of peripheral tissue and systemic energy state. To this end, the liver was first proposed to regulate food intake through glucose sensing[39]. Numerous subsequent publications demonstrated increased food intake in association with vagal transmission [40–43] of chemically reduced hepatic fatty acid oxidation [44–52] and/or depleted hepatic ATP levels in rodents [53–56]. Importantly, oral delivery of the fatty acid oxidation inhibitor etomoxir to human subjects stimulated acute food intake [57]. However, the high potential for off-site action of intraperitoneal delivered chemical inhibitors and use of vagotomy reduced the rigor of these findings. The use of direct genetic manipulations of hepatic energy state on appetite control will be necessary to elucidate the presence of a liver interoceptive sensing mechanism.

Previously, we have reported increased short-term diet-induced weight gain and greater high-fat diet food intake in rats with reduced hepatic MEM [58–60]. However, the hepatic mitochondrial phenotype is present as part of the models differences in intrinsic aerobic fitness based on selective breeding [61], complicating determination of any mechanism(s) responsible for the differences in energy homeostasis. More recently, we observed increased short-term diet-induced weight gain, high-fat diet food intake, and within-meal consumption in a mouse model of reduced liver MEM due to liver-specific reductions in the expression of the mitochondrial co-transcriptional regulator, PGC1a (LPGC1a+/-) [62]. However, it was still unclear whether reduced liver MEM in these mice negatively impacted tissue ATP homeostasis and meal termination.

Combining these data with the previous pharmacological findings, we hypothesize that inhibition of food intake via nutrient-induced, GI-derived satiation signals is decreased under conditions of impaired hepatic ATP homeostasis. To test this hypothesis, we again used the liver-specific, PGC1a heterozygous mouse model. We and others have shown that LPGC1a+/- mice have reduced expression of β-oxidation genes, impaired mitochondrial complete fatty acid oxidation, and diminished mitochondrial respiratory capacity [62; 63]. Herein, we assessed whether the LPGC1a+/- mouse has impaired food intake inhibition in response various nutritional challenges to finally establish a direct link between hepatic MEM and satiation.

## 2. Methods

### 2.1. Ethical Approval

The animal protocol was approved by the Institutional Animal Care and Use Committee at the University of Kansas Medical Center. All experiments were carried out in accordance with the *Guide for the Care and Use of Laboratory Animals* published by the US National Institutes of Health (NIH guide, 8th edn, 2011). Male liver-specific PGC1a heterozygous (LP) mice were produced as previously described [62; 63]. Briefly, male C57Bl6/J mice (#000664, Jackson Laboratory, Bar Harbor, ME, USA) were mated with female PGC1a homozygous floxed mice (#009666, Jackson Laboratory, Bar Harbor, ME, USA) to generate heterozygous PGC1a flox mice. Female PGC1a fl/+ mice were bred to albumin-cre recombinase mice (#003574, Jackson Laboratory, Bar Harbor, ME, USA) to generate liver-specific PGC1a heterozygous and wildtype (WT) littermates. Mice were housed at 28⁰C on 12/12 reverse light cycle (dark 10:00 – 20:00), with *ad lib* access to water and normal chow.

### 2.2. Genotyping

Mouse and liver-specific genotypes were confirmed as previously described [63]. The primers used are listed below.

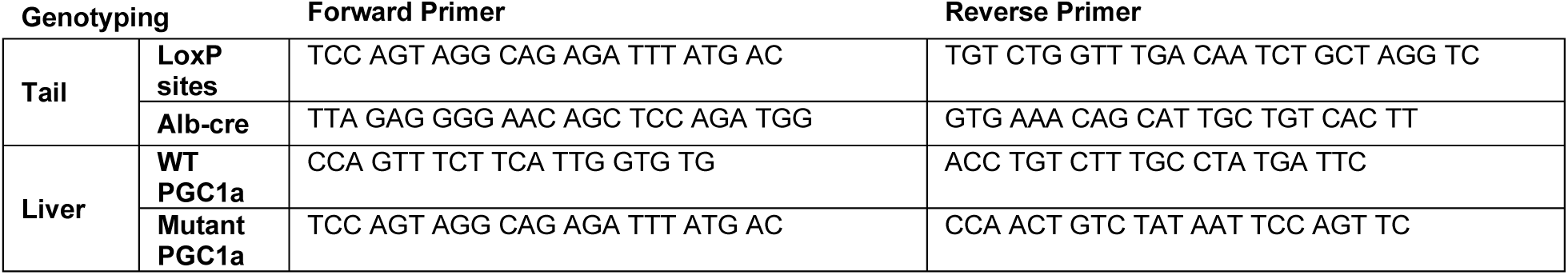
Primer table.

### 2.3. Food Intake Regulation

LP and WT mice were acclimated to individual housing and oral gavage in BioDAQ cages (Research Diets Inc., New Brunswick, NJ) for 3 weeks prior to the initiation of the food intake regulation studies. On experimental days (Monday & Thursday), mice were food restricted at 0800, underwent gavage at 0930, and food access and data collection began at 1000. Sham gavage experiments were performed periodically to assess basal feeding throughout the study (n-4 sham). Additional, basal food intake data was collected during the non-experimental days (n=9). Basal food intake data presented represents the combined average of each mouse across all sham treatments and non-experimental days collected (**Figure 1A**). Ensure powder (Abbot Nutritional Products, Abbott Park, IL) was resuspended in MilliQ water to produce a 5 kcal/mL solution, and dosed at 5, 10, and 15 mL/kg. This resulted in the delivery of 1.5 kcal at the 10 mL/kg dose, which represents about ∼15% of the kcals consumed in 24hr or the approximate energy intake during the first 2 hrs of the dark cycle. Methylcellulose (M0512, Sigma Aldrich, St. Louis, MO) was dissolved in normal saline at 1% & 2% by weight and delivered at 10 ml/kg. Antagonists of peripheral satiation receptors 5-HT3R (ondansetron, 5 mg/kg, saline, PHR1141, Sigma Aldrich, St. Louis, MO), CCK-1R (lorglumide, 0.3 mg/kg, saline, 17555, Caymen Chemicals, Ann Arbor, MI), and GLP1R (exendin-9, 100 ug/kg, saline, 4017799, Bachem, Torrance, CA) were delivered IP, 30 minutes before the initiation of oral gavage of 10 mL/kg Ensure. Intralipid (Baxter, Deerfield, IL), 40% glucose (D16, Thermo Fisher Scientific, Waltham, MA), and Proteinex (Llorens Pharmaceutical, Miami, FL) were delivered via oral gavage at 10 and 15 mL/kg to assess specific macronutrient oral pre-load inhibition of food intake. This results in the delivery of 0.6 kcal of lipid and protein, and 0.48 kcal of glucose at the 10 mL/kg dose. For fasting-induced acute food intake experiments, the BioDAQ gates were closed at 1600 resulting in an 18hr fast, with food intake measurements beginning at 1000. BioDAQ data analysis was binned at 0.5 hr or 1 hr intervals using BioDAQ Data Viewer (ver. 2.3.14, BioDAQ, Research Diets Inc. New Brunswick, NJ), with inter-meal interval and minimum meal mass set at 90 seconds and 0.05 grams, respectively.

**Figure 1.**
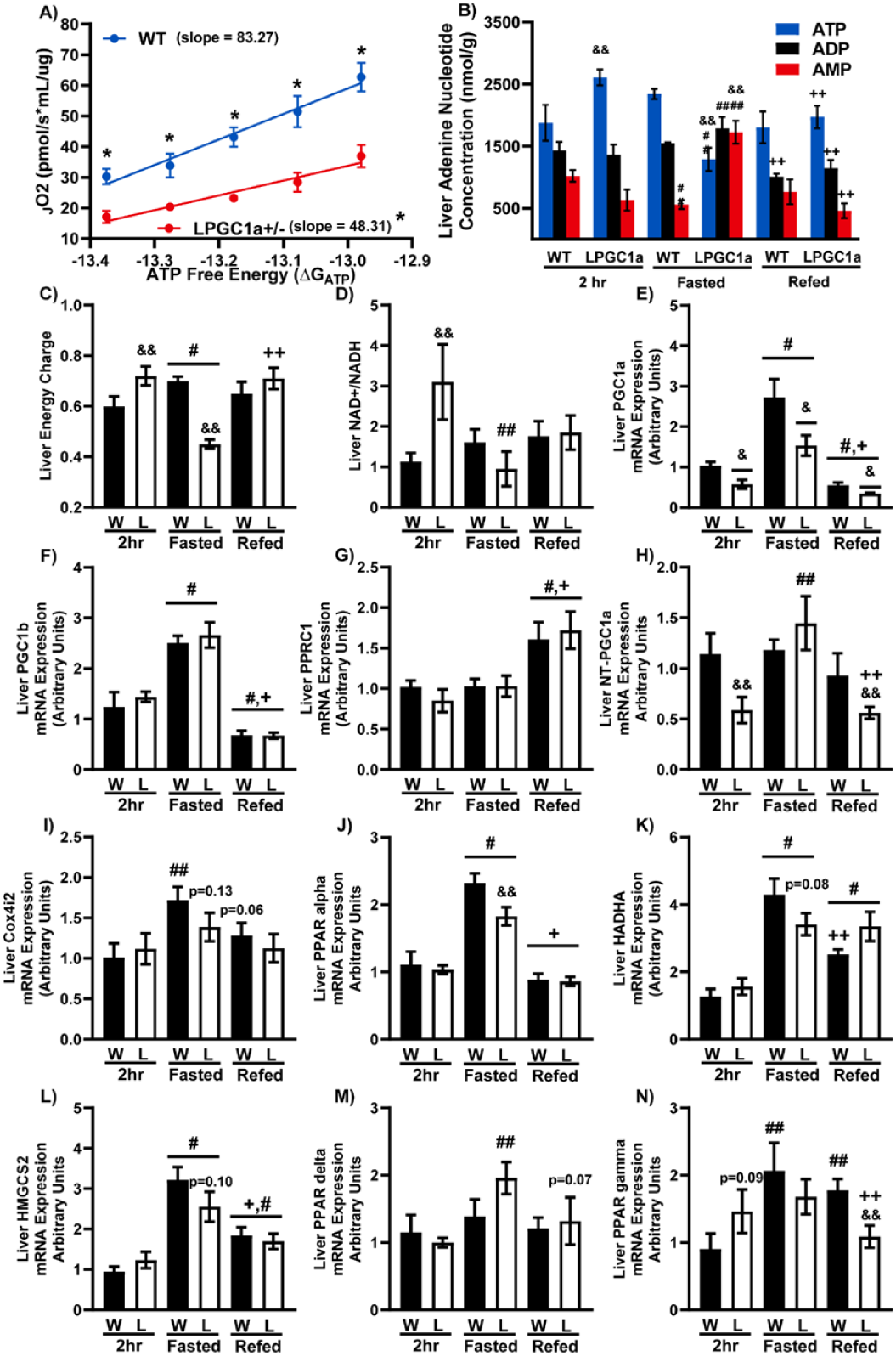
LPGC1a+/- Mice Have Reduced ATP Homeostasis During Fasting. A) Respiration of isolated liver mitochondria during changes in ATP free energy (ΔG ATP) via creatine kinase clamp. B) Liver ATP, ADP, & AMP concentration (nmol/g) in mice after 2hr food withdrawal, 18 hr fast, or 4 hr refeeding. Impact of fasting/re-feeding on the liver was assessed as C) liver energy charge (ATP + (0.5 x ADP))/(ATP+ADP+AMP), D) NAD+/NADH, and mRNA expression of E) PGC1a, F) PGC1b, G) PPRC1, H) NT-PGC1a, I) Cox4i2, J) PPAR alpha, K) HADHA, L) HMGCS2, M) PPAR delta, and N) PPAR gamma in 2hr food withdrawn, 18hr fasted, or 4 hr refed mice. Data are represented as mean ± SEM. n=3-8 biological replicates for each genotype for adenine nucleotide concentration. n=8-10 biological replicates for each genotype for all other data. # main effect fasting or refeeding compared to 2hr, & main effect of genotype, + main effect of refed vs fasted by two-way ANOVA. *p<0.05 between genotypes by Student’s t-test (A). # main effect vs 2hr, + main effect refed vs fasted, & main effect LPGC1a+/- vs WT. ## fasting versus 2 hr within genotype, && LPGC1a+/- versus wildtype within group, and ++ refed versus fasting within genotype pairwise comparisons were performed using Fishers LSD. See also Figure S5 for liver mRNA expression of mitochondrial in 2hr food withdrawn, fasted, and 4 hr refed mice.

### 2.4. Plasma GI-derived Satiation Signal Concentration

Following a 2 hr food restriction mice received a 5 mL/kg oral gavage of Ensure, as described above. Blood was collected 90 minutes following the oral nutrient gavage in anesthetized mice. Blood was added to EGTA coated tubes, plasma was collected following centrifugation (4°C, 8,000 x g, 10 min), and stored at -80°C. Plasma 5-HT (LS-F40041, LS Bio, Shirley, MA), CCK (EIAM-CCK, RayBiotech, Norcross, GA), and GLP-1 (LS-F23033, LS Bio, Shirley, MA) concentration was determined as described by the manufacturer.

### 2.5. Liver Mitochondria Isolation

Liver mitochondria were isolated as previously described[62]. Briefly, ∼ 1 g of liver was homogenized (glass-on-teflon) in 8 mL of ice-cold mitochondrial isolation buffer (220 mM mannitol, 70 mM sucrose, 10 mM Tris, 1 mM EDTA, pH adjusted to 7.4 with KOH). The homogenate was centrifuged (4°C, 10 min, 1500 g), the supernatant was transferred to a round bottom tube, and centrifuged (4°C, 10 min, 8000 x g). The pellet was resuspended in 6 mL of ice-cold mitochondrial isolation buffer using a glass-on-glass homogenizer, and centrifuged again (4°C, 10 min, 6000 x g). The pellet was resuspended in 4 mL of isolation buffer containing 0.1% BSA and centrifuged (4°C, 10 min, 4000 x g). This final pellet was resuspended in ∼0.75 mL of mitochondrial respiration buffer (0.5 mM EGTA, 3 mM MgCl2, 60 mM KMES, 20 mM glucose, 10mM KH2PO4, 20 mM HEPES, 110 mM sucrose, 0.1% BSA, 0.0625 mM free CoA, and 2.5 mM carnitine, pH∼7.4). The protein concentration for both suspensions was determined by BCA assay.

### 2.6. Liver Mitochondrial Creatine Kinase Clamp

Change in respiratory rate of isolated liver mitochondria during changes in ΔG ATP were performed as previously described with minor changes [64; 65]. Briefly, 2 mM malate, isolated liver mitochondria, 10 μM palmitoyl-CoA, 10 μM palmitoyl-carnitine, 5 mM ATP, 5 mM creatine, and 20 U/mL creatine kinase were added to Oroboros chambers containing mitochondrial respiration buffer. ADP-dependent respiration was initiated by the addition of 1 mM PCr, and sequential additions of PCr to 3, 6, 9, 12, 15, 18, 21, 24, 27, 30 mM reduced respiration toward baseline. The free energy of ATP for each PCr concentration was calculated as described [64]. The linear respiration data from 9 mM to 21 mM PCr was used and the conductance represented as the slope of the line. The data is oriented to represent an simulated increase in energy demand produced by the clamp.

### 2.7. Citrate Synthase Activity

Citrate was determined as previously described[66]. Briefly, frozen liver was homogenized in 0.175 mM KCl, 2.0 mM EDTA (pH = 7.4) and centrifuged at 2,000 x g (4°C) for 5 minutes. Supernatants were freeze/thawed three times prior to initiation of assay. Aliquots of tissue homogenate were diluted in reaction buffer (100 mM Tris, 1.0 mM DTNB, 10 mM oxaloacetate, pH∼8.3). The reaction was initiated by the addition of 3 mM acetyl-CoA, and kinetic reads were taken at 412 nm every minute for 7 minutes (37°C). Values were normalized to BCA of the homogenate.

### 2.8. Fasting/Re-feeding

Mice had food withdrawn for 2hrs or overnight (18hrs, 1600 – 1000). Re-feed mice were allowed access to normal chow for 4 hrs following the fast. Mice were anesthetized with 100 mg/kg phenobarbital, and a section of the liver was clamp frozen. The brain was frozen in 2-methyl-butane and both tissues were stored at -80°C.

### 2.9. Liver Adenine Nucleotide Pool

Measurement of liver adenine nucleotide pool was performed by the University of Minnesota Metabolomics Core. Clamp frozen liver sections were weighed and homogenized using zirconium beads in 0.4 M perchloric acid, 0.5 mM EGTA extraction solution containing [^13^C_10_, ^15^N_5_]ATP sodium salt (100 mM), [^13^C_10_, ^15^N_5_]AMP sodium salt (100 mM), [1,2-^13^C_2_]acetyl-CoA lithium salt (5 mM), and [1,2,3-^13^C_3_] malonyl-CoA lithium salt (5 mM). After incubation on ice for 10 min, samples were centrifuged at 15,000xg for 15 min at 4⁰C. The resulting supernatants were neutralized with freshly prepared 0.5 M K_2_CO_3_, vortexed, and centrifuged at 15,000xg for 30 min at 4⁰C. Final extracts were then analyzed by LC-MS/MS as previously described [67; 68], with modifications. Energy charge was calculated as (ATP + (0.5 x ADP))/(ATP+ADP+AMP).

### 2.10. Liver & Hypothalamus RNA Expression

Liver RNA was isolated from ∼ 25 mg of liver tissue using an RNeasy Plus Mini Kit (Qiagen, Valencia, CA, USA) and hypothalamus RNA was isolated using the standard Triazol procedure. The cDNA for both tissues was produced using the ImProm-II RT system (Promega, Madison, WI, USA). RT-PCR was performed using a Bio-Rad CFX Connect Real-Time System (Bio-Rad, Hercules, CA, USA) and SYBR Green. The SYBR green primers are listed below. Gene specific values for liver were normalized to relative cyclophilin B (*ppib*) mRNA expression values and hypothalamus gene specific values were normalized to Tryptophan 5-Monooxygenase Activation Protein Zeta (*ywhaz*) mRNA expression values.

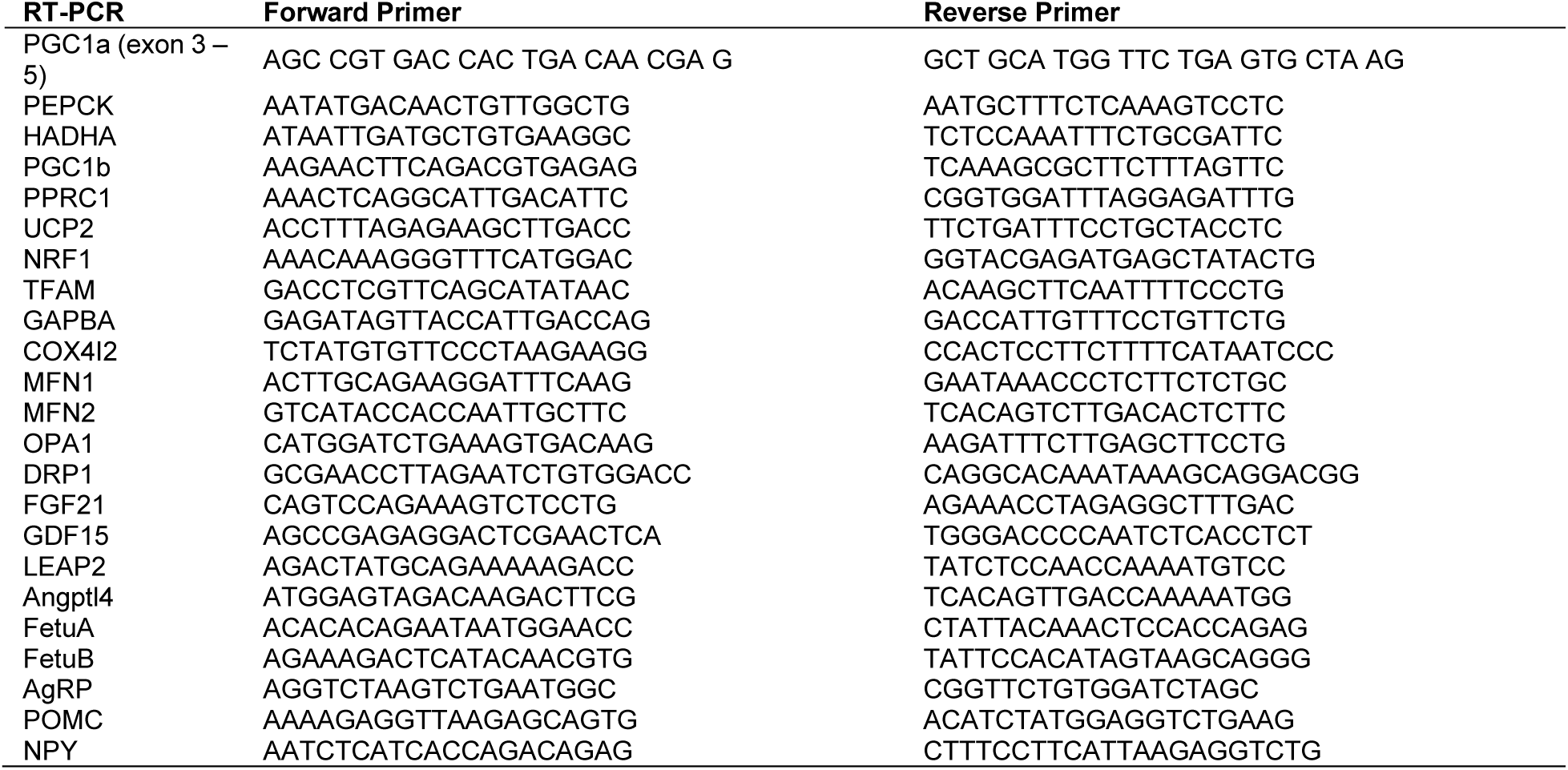

### 2.11. Liver Glycogen Concentration

Liver glycogen was determined as previously described [69]. Briefly, ∼15 to 30 mg of skeletal muscle was solubilized in 0.5 mL of 1 N HCl in boiling water for 1.5 to 2.0 hours. Samples were neutralized by adding 1.5 mL of 0.67 M NaOH. Free glycosyl units were determined spectrophotometrically using a glucose oxidase kit.

### 2.12. Fluorescent *In Situ* Hybridization

Four-hour fasted wildtype and LPGC1a+/- mice were transcardially perfused with 0.1 M PBS followed by 4% paraformaldehyde in 0.1 M PBS on ice. Brains were removed, post-fixed in 4% paraformaldehyde for 24h, then stored in 20% sucrose in 0.1 M PBS at 4 °C until sunk. Coronal hypothalamic sections (20 μm) were sliced on a cryostat, dry mounted onto slides, and stored at -80°C for future fluorescent in situ hybridization (FISH) processing. FISH to quantify mRNA expression of AgRP and POMC neuropeptides was performed using the RNAscope Multiplex Fluorescent V2 kit (323110, Advanced Cell Diagnostics, Inc.) according to manufacturer’s instructions. The probes used were: mmPOMC (314081, Advanced Cell Diagnostics, Inc.) and mmAgRP (400711-C2, Advanced Cell Diagnostics, Inc.). Upon completion of the FISH protocol, slides were then coverslipped with Fluoromount G (0100-01, SouthernBiotech). Slides were visualized using fluorescence microscopy (BZ-X810, Keyence) and image analysis to quantify fluorescent signal of POMC and AgRP at weak, moderate, and strong intensity levels was performed using the HALO® Area Quantification FL module and expressed as total fluorescent area (µm^2^) .

### 2.13. Statistical Analysis

Data are presented as means and standard error. The two-standard deviation test was utilized to test for outliers within group. Food intake, mitochondrial respiration, and *in situ* hybridization data was assessed using the Student’s t-test between genotypes within time. Fasting/re-feeding data was assessed by two-way ANOVA. Pairwise comparisons were performed using Fisher’s LSD. Main effects are only designated only when all pairwise comparisons were significant.

## 3. Results

### 3.1. LPGC1a+/- Mice Have Decreased Liver Mitochondrial Respiration in Response to Increasing Energy Demand and Impaired ATP Homeostasis During Fasting

The liver has been proposed to regulate food intake by serving as a interoceptive peripheral energy sensor through transmission of hepatic energy state via the vagus nerve [70]. While we have previously observed reduced maximal respiration and fatty acid oxidation in isolated liver mitochondria from male LPGC1a+/- compared to WT mice [62], these measures are conducted under non-physiological conditions and do not assess maintenance of cellular energy homeostasis. To assess respiration under physiologically relevant energy demand and to measure the capacity of the mitochondria to increase respiration in response to increased ATP free energy states (conductance) we used the creatine kinase clamp technique in isolated liver mitochondria following a 4 hr food withdrawal [64; 71]. Not only do isolated liver mitochondria from male LPGC1a+/- mice have reduced respiration of fatty acids at all ATP free energy states tested (**Figure 1A**), they also have reduced capacity to respond to increasing energy demand with increased respiration (slope of line across ATP free energy states). This suggests that the male LPGC1a+/- mice will have impaired capacity to maintain hepatic energy status during systemic energy challenges. Subsequently, we used overnight fasting/re-feeding as systemic energy challenge to measure the adenine nucleotide pool of WT and LPGC1a+/- mice (**Figure 1B**). LPGC1a+/- mice were unable to maintain liver ATP levels following the overnight fast, and had an associated increase in AMP levels. After 4 hours of re-feeding this loss of liver energy homeostasis was ameliorated. Additionally, representing the differences in adenine nucleotides as liver energy charge [i.e., (ATP + (0.5 x ADP))/(ATP+ADP+AMP), (**Figure 1C**)], and NAD^+^/NADH ratio (**Figure 1D**) highlights the impaired in maintenance of the hepatic energy state in fasted male LPGC1a +/- mice. The differences in ATP and AMP during fasting were not due to changes in the total adenine nucleotide phosphate pool, which was not different between genotypes at the three timepoints (**Figure S1A**). Interestingly, the total nicotinamide adenine nucleotide pool was increased in male LPGC1a+/- liver during fasting (**Figure S1B**), due to a large increase in NADH concentration (**Figure S1D**). The reductive stress generated by this increased NADH relative to NAD+ is commonly observed in Metabolic Dysfunction-associated Steatotic Liver Disease, and occurs due to decreased capacity of the electron transport system to accept electrons from NADH and FADH_2_ [72]. Importantly, both genotypes experienced a similar level of energy pressure upon the liver as witnessed by the lack of difference between genotypes at all time points for glycogen and gluconeogenic marker PEPCK mRNA expression (**Figure S2A & B**).

The inability to maintain hepatic energy state in LPGC1a+/- mice could be partially due to impaired mitohormesis [73], the maintenance of a highly functioning pool of mitochondria during varying physiological conditions is key to appropriate energy stress response. While we [62; 74] and others [63] have observed reduced mitochondrial respiration, fatty acid oxidation, mitophagy, and beta-oxidation gene expression in the male LPGC1a+/- mouse, the defect(s) that result in impaired liver mitochondrial energy metabolism are not completely known. As expected, liver PGC1a expression was ∼50% reduced in LPGC1a+/- mice compared to WT during all phases of the fasting/re-feeding experiment (**Figure 1E**). Importantly, no genotype specific gene expression differences in PGC family members, PGC1b and PPRC1 were observed during the fasting/re-feeding experiments (**Figures 1F & 1G**). Demonstrating that heterozygous expression of PGC1a in hepatocytes did not elicit a compensatory expression in these PGC1 family genes. However, mRNA expression of the shorter alternative 3’ splicing variant, N-terminal PGC1a (NT-PGC1a), was approximately 50% lower under basal and refed conditions in LPGC1a+/- mice compared to WT (**Figure 1H**). During fasting no difference in NT-PGC1a expression was observed between genotypes, however, fasted LPGC1a+/- mice greater NT-PGC1a expression compared to basal and refed. Liver NT-PGC1a expression can be induced by fasting [75], it is unclear why only the LPGC1a+/- show a fasting response. However, fasting-induced liver NT-PGC1a expression may explain the maintenance of fasting response for many mitochondrial and gluconeogenic [76] genes in LPGC1a+/- mice. PGC1a regulates mitohormesis partially through transcriptional co-regulation of transcriptional pathways regulating mitochondrial biogenesis and oxidative metabolism [73; 77]. While nuclear respiratory factor-1 (NRF1) [78] showed reduced liver gene expression under basal and fasted conditions in LPGC1a+/- mice, no genotype effects were observed for nuclear respiratory factor-2 (GABPA) [78] or mitochondrial transcription factor-a (TFAM) [79] (**Figure S2D, E & F**, respectively). Cumulative changes in the activity of these transcriptional pathways regulate markers of mitochondrial content. Liver citrate synthase activity was not affected by genotype or fasting/refeeding (**Figure S2C**). However, liver expression of the Cox IV nuclear subunit, Cox4i2, was increased by fasting WT and tended to be greater compared than LPGC1a (**Figure 1I**). Also, fasting did not significantly increase LPGC1a+/- Cox4i2 expression compared to basal. This reduction in expression of an electron transport system gene in LPGC1a+/- mice suggests that the increases in liver NADH (**Figure S1D**) and reduced NAD+/NADH (**Figure 1D**) during fasting are due to decreased capacity to accept and shuttle electrons. Further, the increased hepatic NADH during fasting could generate the observed increased liver acetyl-CoA (**Figures S1E**), decreased succinyl-CoA (**Figure S1F**), and the decreased mitochondrial TCA cycle flux [62] in LPGC1a+/- mice putatively through product inhibition of TCA cycle dehydrogenases [80].

PGC1a was discovered as a transcriptional co-regulator of PPARγ in brown adipose [81], and has been shown to regulate both oxidative metabolism pathways through regulation of PPARα and δ [77]. Fasting increased PPARα mRNA expression in both genotypes, however, LPGC1a+/- mice had a smaller increase (**Figure 1J**). This fasting increase in PPARα expression was mirrored by the expression of the enzymes involved in mitochondrial (HADHA) and peroxisomal (ACOX1) β-oxidation (**Figure 1K & S2M**, respectively), the rate limiting enzyme in the mitochondrial uptake of long-chain acyl-CoA (CPT1a, **Figure S2G**), mitochondrial uncoupling protein 2 (UCP2, **Figure S2L**), and the rate limiting enzyme in ketone synthesis (HMGCS2, **Figure 1L**) over basal in both genotypes. However, only HADHA and HMGCS2 tended to have higher expression in WT compared to LPGC1a during fasting. While LPGC1a+/- liver mitochondrial β-oxidation may be reduced during fasting, the lack of difference in CPT1a expression and the reduced malonyl-CoA concentration (**Figure S1G**) suggest entry of fatty acids into the mitochondria is not reduced. Interestingly, liver PPARδ was not different across the stimuli in WT mice, but was increased by fasting in the male LPGC1a+/- mice (**Figure 1M**). Also, liver PPARγ expression tended to be higher in LPGC1a+/- under basal conditions (**Figure 1N**) but was only increased by fasting in WT mice. While PGC1a controls mitohormesis through regulation of mitochondrial quality control pathways [82], no genotype differences in basal or fasted expression of genes involved in mitochondrial fusion or fission was observed (**Figure S2H – K**). Finally, PGC1a can regulate hepatocyte mitochondria through the activity of transcription factors hepatocyte nuclear respiratory factor 4a (HNF4a [83]) and forkhead box O1 (FOXO1 [84]). Again, no differences were observed between genotypes in the liver expression of these transcription factors under basal, fasting, or refed conditions (**Figure S2O & P**, respectively).

Recent work demonstrates that changes in the hepatocyte membrane potential are part of a powerful signaling pathway linking peripheral energy metabolism and central regulation of metabolism, including food intake [85; 86]. The primary component of cellular membrane potential development and maintenance is the action of the Na/K-ATPase. In hepatocytes, the Na/K-ATPase is primarily comprised of the ATP1a1 catalytic subunit, either ATP1b1 or ATP1b3 subunit, and the FXYD1 regulatory subunit (reviewed [87; 88]). The action of the hepatocyte Na/K-ATPase represents ∼30% of cellular oxygen consumption [89], making it a large consumer of cellular ATP. Interestingly, fasting generates increased liver mRNA expression of ATP1a1, ATP1b1, and FXYD1 in male WT mice (**Figure S3A - D**). However, LPGC1a+/- mice have less of an increase in ATP1a1, and no fasting-mediated changes in FXYD1. Also, the plasma membrane calcium ATPase, ATP2b1, is increased by fasting in both genotypes, but to a lesser extent in LPGC1a+/- mice (**Figure S3G**). While not representative of the activity of these ion pumps, these data suggest impaired capacity to maintain membrane potential, particularly during energy stress, in the male LPGC1a+/- mouse. Future work will focus on the role of impaired hepatocyte mitochondrial ATP homeostasis on the function of the Na/K-ATPase system and maintenance of the cellular membrane potential.

Together these data showing PGC1a expression plays a unique role in the regulation of liver mitochondrial transcriptional and energy metabolism pathways. Further, it demonstrates that the livers of male LPGC1a+/- mice lack the capacity to respond to physiologically relevant energy stress to maintain tissue ATP levels.

### 3.2. Reduced Liver-Specific PGC1a Results in Impaired Satiation

Previously, we have observed that male LPGC1a+/- mice have increased high-fat/high-sucrose diet food intake and weight gain compared to wildtype (WT) littermates, however, no differences were observed in low-fat diet food intake and body weight [62]. Here, we again see no difference in cumulative chow intake between LPGC1a+/- and WT mice in the first three hours of the dark cycle (**Figure 2A**). However, upon examination of meal microstructure, feeding duration per 1 hour period was significantly increased (**Figure 2B**) and there was a tendency for greater average meal size per 1 hr period (**Figure 2C**) in male LPGC1a+/- mice compared to wildtype. To offset this increase in meal size and duration the LPGC1a+/- mice on average consumed significantly fewer meals per 1 hr period (**Figure 2D**). These alterations in feeding patterns were particularly evident at the 2 hr time point. These data suggest an association between reduced hepatic MEM and impaired within meal satiation, and further suggest that these underlying impairments in within meal satiation require increased dietary energy density to elicit greater food intake.

**Figure 2.**
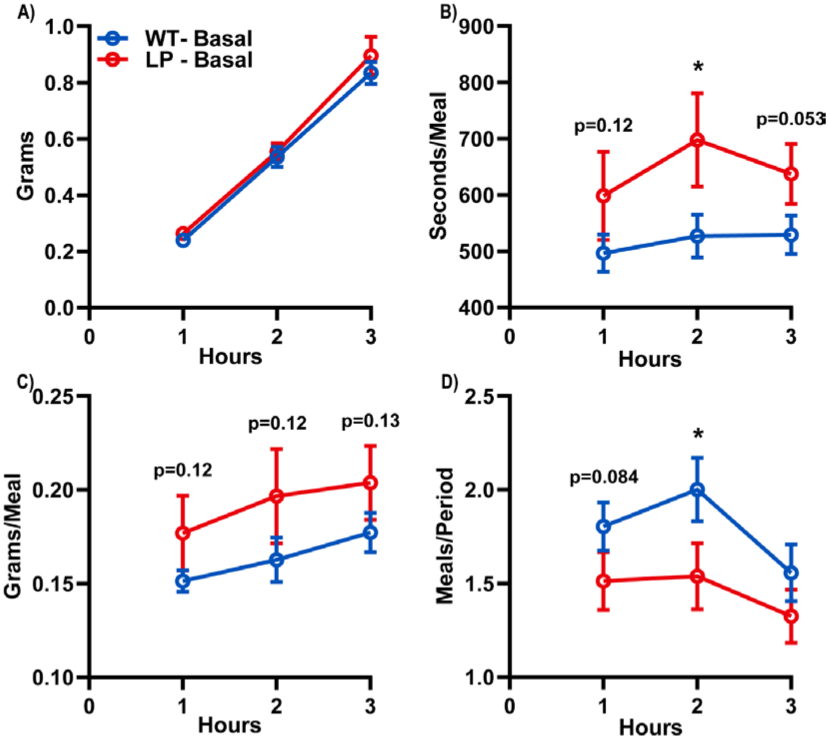
Reduced Liver-Specific PGC1a Results in Underlying Differences in Feeding Behavior. A) Acute basal food intake at the beginning of the dark cycle following 2hr food withdrawal. B) Average number of meals, C) grams per meal, and D) length of each meal during each 1hr period. Data are represented as mean ± SEM. n = 13 technical replicates and n= 7 – 9 biological replicates for each genotype. *p<0.05 between genotypes within time by Student’s t test.

### 3.3. LPGC1a+/- Mice are Less Sated by a Mixed Macronutrient Gastric Preload

Previous work suggesting a role of liver MEM in food intake regulation were confounded by the lack of target tissue specificity due to the use of peripherally delivered nutrients [44; 90-93], and IP delivery of molecules to acutely inhibit liver oxidative metabolism [40; 45; 48; 52] or lower hepatocyte ATP [49; 54-56]. The hepatocyte-specific nature of the reduction in liver MEM of the LPGC1a+/- mouse model allows for more rigorous investigation of the role of liver energy metabolism on food intake regulation. Based on the previously observed increase in high-fat diet-induced food intake [62] and the underlying difference in meal pattern structure (**Figure 2**), we next assessed whether reduced liver MEM impaired the food intake suppressive effects of oral mixed nutrient preloads. Mice were presented with chow following the oral gavage and meal patterning was assessed over the next 3 hours. While oral gavage of 5, 10, and 15 mL/kg Ensure (5 kcal/mL) resulted in dose-dependent inhibition of chow intake in male WT and LPGC1a+/- mice relative to sham gavage **(Figure 3A-C, M-O)**, male LPGC1a+/- mice had reduced suppression of food intake at all time points and doses compared to WT. As in **Figure 2**, this was due to increased meal duration (**Figure 3E-G**) and size (**Figure 3I-K**) at various time points during the different oral preload doses. Interestingly, unlike **Figure 1**, no differences were observed in the number of meals between genotypes except at the highest dose (**Figure S4A-C**). The inhibition of food intake by oral gavage of a mixed meal is due to satiation signals elicited by the nutrients and distention of the GI tract. To verify that the impaired food intake inhibition of the male LPGC1a+/- mice was due to GI-derived nutrient signals and not dysregulated sensing of mechanical distention of the GI tract we administered non-nutritive methylcellulose oral pre-loads. Oral gavage of a 1% methylcellulose solution inhibited food intake to a greater extent in male WT mice compared to LPGC1a+/- (**Figure 3D**). However, this was not due to significant differences in meal duration (**Figure 3H**), size (**Figure 3L**), or number (**Figure S4D**) as was observed with mixed nutrient. Further, a dose dependence of non-nutritive mechanical food intake inhibition was observed as an oral preload of 2% methylcellulose resulted in the same food intake inhibition in both genotypes (**Figure S4E-H**). These data demonstrate that mice with reduced liver MEM have impaired meal termination in response to GI-derived satiation signals initiated by oral nutrients.

**Figure 3.**
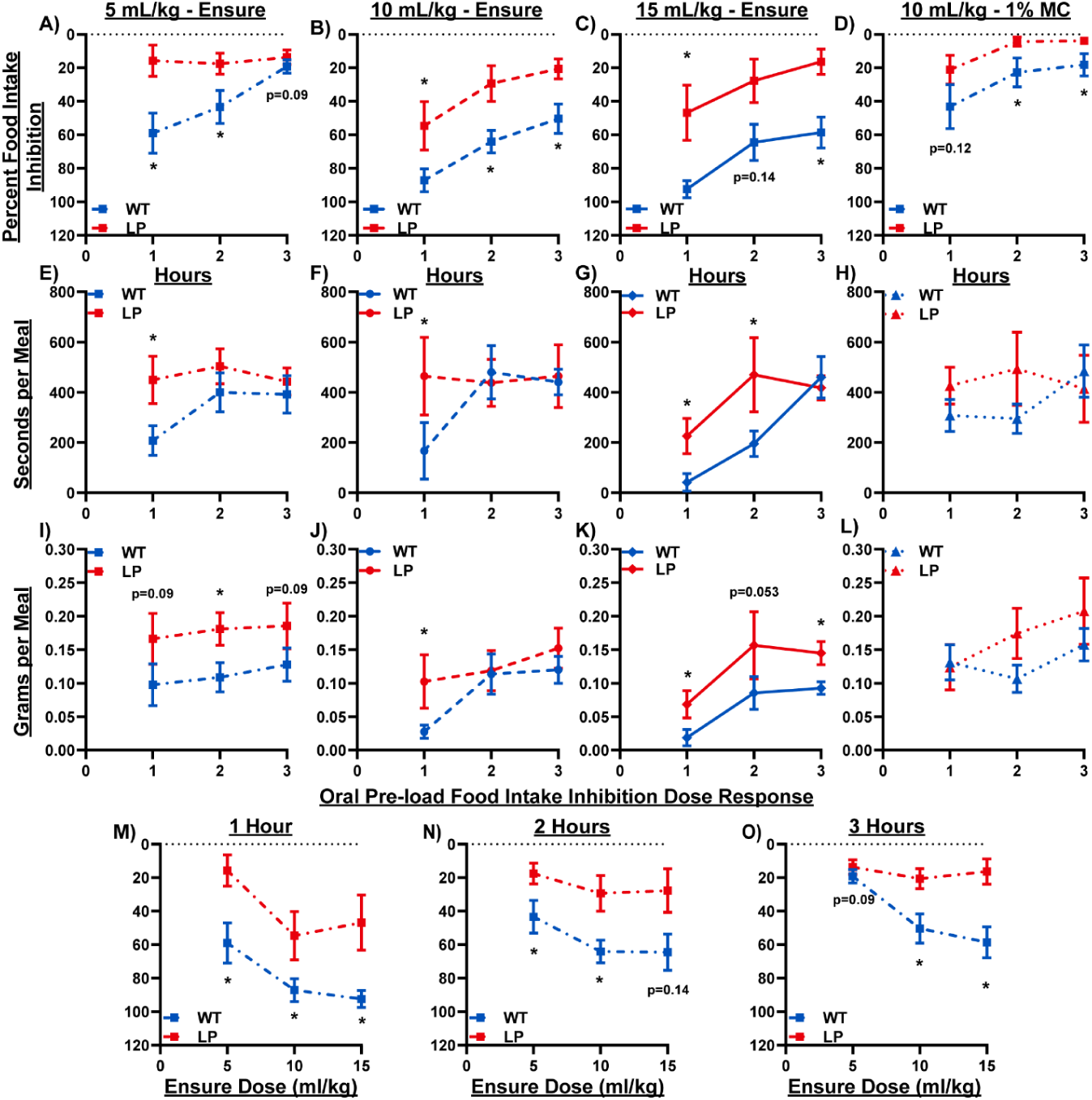
Dose Dependent Impairment in Satiation Following Mixed Macronutrient Oral-Preloads in LPGC1a+/- Mice. Percent inhibition of food intake following oral pre-load of Ensure® at A) 5, B) 10, & C) 15 mL/kg and D) 10 mL/kg 1% methylcellulose. Average duration of meal per 1 hr period following mixed nutrient E) 5, F) 10, & G) 15 mL/kg, and H)10 m/kg methylcellulose oral pre-load. Average meal size period 1 hr period following I) 5, J) 10, & K) 15 mL/kg nutrient, or L) 10 mL/kg non-nutritive oral pre-load. Ensure® oral pre-load food intake inhibition dose response at M) 1 hr, N) 2 hr, & O) 3 hr. Data are represented as mean ± SEM. n= 7 – 9 biological replicates for each genotype. *p < 0.05 between genotypes within time by Student’s t test. See also Figure S2 for meals per period data across the 3 oral pre-load doses.

### 3.4. Decreased Satiation Response in LPGC1a+/- Mice Following Gastric Delivery of Individual Macronutrients

GI delivery of carbohydrate, protein, and lipids produces acute suppression of food intake in mice and humans (reviewed [6; 17]). The differences in food intake and feeding behavior following oral gavage of a mixed nutrient observed in Figure 2 could be due to impairments in the capacity of individual macronutrients to elicit a satiation response. Therefore, we determined whether acute food intake inhibition following oral gavage of individual macronutrients was reduced in male LPGC1a+/- mice. Percent inhibition of food intake 1 hr following oral gavage of glucose was lower in LPGC1a+/- mice **(Figure 4A**) and tended to be lower following lipid and protein delivery **(Figure 4B & C)**. Only protein continued to show consistent reduction of food intake inhibition in LPGC1a+/- throughout the 3-hour data collection. No differences were present at 2 and 3 hours after lipid delivery, and oral gavage of glucose increased food intake inhibition in LPGC1a+/- mice at 2 and 3 hours compared to WT. Overall, feeding behavior differences were more subtle for the individual macronutrients.

**Figure 4.**
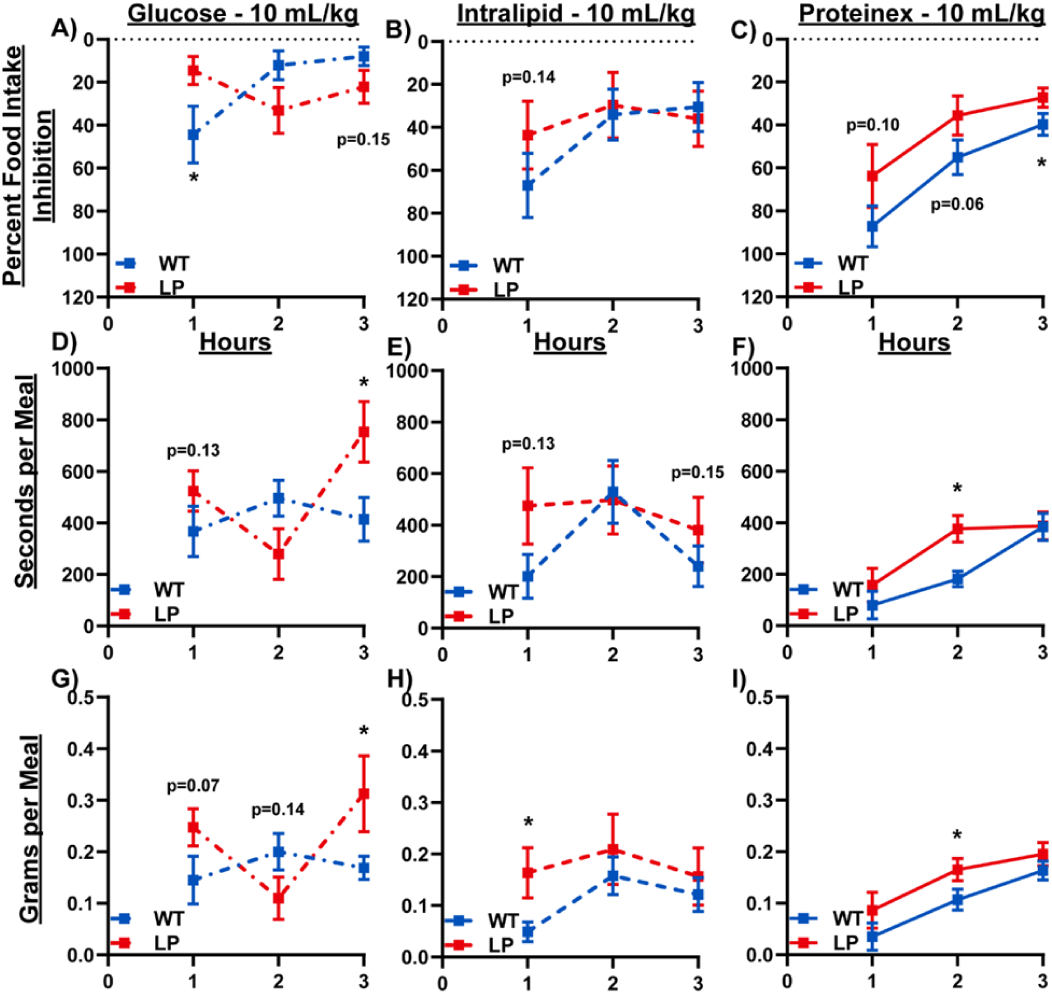
Decreased Satiation Response in LPGC1a+/- Mice Following Oral Delivery of Individual Macronutrients. Percent inhibition of food intake following oral pre-load of individual macronutrients at 10 mL/kg: A) 40% Glucose, B) Intralipid®, & C) Proteinex®. Average duration of meal per 1 hr period following D) 40% Glucose, E) Intralipid®, & F) Proteinex® individual macronutrient oral pre-load. Average meal size per 1 hr period following G) 40% Glucose, H) Intralipid®, & I) Proteinex® nutrient oral pre-load. Data are represented as mean ± SEM. n= 7 – 9 biological replicates for each genotype. *p < 0.05 between genotypes within time by Student’s t test. See also Figure S3 for meals per period data across the 3 individual macronutrient oral pre-loads.

Glucose oral pre-load showed time dependent difference between male WT and LPGC1a+/- mice (**Figure 4D & G**). Oral lipid tended to increase meal duration at 1 & 3 hr (**Figure 4E**) in LPGC1a+/-, and increased meal size only at 1hr (**Figure 4H**). Following protein oral pre-load, greater meal duration and size were only observed in LPGC1a+/- mice at 2hrs (**Figures 4F & I**). The number of meals following oral gavage of the three macronutrients was not different, except glucose with LPGC1a+/- mice consuming fewer meals at 2 & 3 hrs (**Figure S5**).

While the individual macronutrient oral pre-loads generated differences in food intake inhibition and feeding patterns, the data were not as striking as observed with the mixed macronutrient. The reduced caloric density of the oral nutrient pre-loads at the 10 mL/kg dose (∼0.6 kcal – lipid & protein, ∼0.48 kcal – glucose compared to ∼1.5 kcal of Ensure) may explain the more subtle differences observed between male LPGC1a+/- and WT littermates. However, these findings suggest that the energy density of the nutrient loads may be involved in eliciting a greater phenotype. This could explain why the only the high-fat diet fed male LPGC1a+/- mice had greater within meal food intake in our previous work [62]. Further, these data demonstrate that the reduced liver MEM in the male LPGC1a+/- mice is associated with reduced food intake inhibition in response to pre-loads of all three individual nutrients tested, suggesting the impaired satiation is not specific to one macronutrient.

### 3.5. Peripheral Satiation Signal Receptor Inhibition Does Not Increase Food Intake Following Oral Pre-Loads in LPGC1a+/- Mice

The above findings demonstrate that male LPGC1a+/- have altered basal feeding behavior and impaired food intake inhibition in response to nutritive and non-nutritive pre-loads. But, from these data it is not possible to discern if a specific satiation signaling pathway is involved. To assess whether function of the peripheral serotonin (5HT), cholesystekinin (CCK-8), and glucagon-like peptide-1 (GLP-1) signaling pathways are impaired in the regulation of food intake inhibition in LPGC1a+/- mice, we delivered selective receptor antagonists 30 minutes prior to a 10 mL/kg Ensure oral pre-load. Intraperitoneal injection of antagonists for 5-HT3R (ondansetron), CCK-1R (lorglumide), and GLP1R (Exendin-9) resulted in significantly greater increases in food intake following the nutritive oral pre-load in male WT compared to LPGC1a+/- mice relative to 10 ml/kg Ensure oral pre-loads alone (**Figures 5A, B, & C**, respectively). 5-HT3R antagonism was able to increase LPGC1a+/- food intake following Ensure, however, CCK-1R and GLP1R blockade had little to no effect on food intake in male LPGC1a+/- mice. Importantly, no difference in serum concentration of 5-HT, CCK-8S, or GLP-1(7-36) were observed between male WT and LPGC1a+/- mice 60 minutes after Ensure oral gavage (**Figure S6A-C**). These findings suggest that the impairment in food intake inhibition observed in LPGC1a+/- mice is due to diminished or modulated vagal afferent satiation signaling rather than decreased release of meal derived GI-satiation signals.

**Figure 5.**
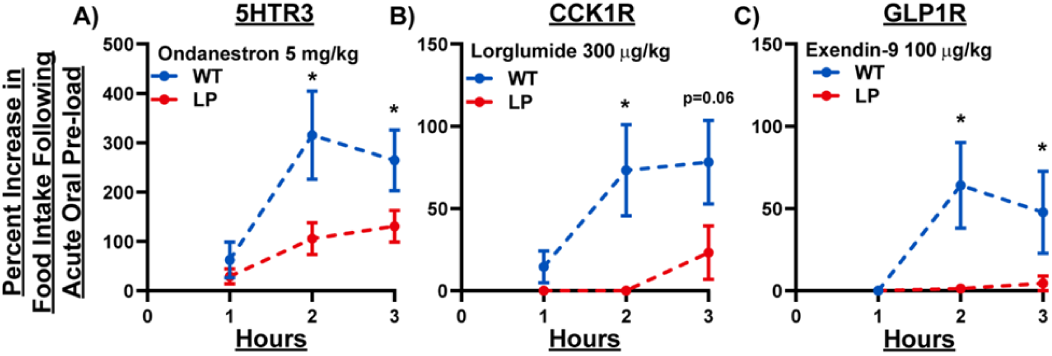
Peripheral Satiation Signal Receptor Inhibition Does Not Impact Food Intake Following Oral Pre-Loads in LPGC1a+/- Mice. Percent increase in chow intake during acute mixed nutrient oral pre-load (10 mL/kg) following intraperitoneal delivery of satiation signal receptor inhibitors A) Ondanestron (5-HT3R antagonist, 5 mg/kg), B) Lorglumide (CCK-1R antagonist, 300 mg/kg), and C) Exendin-9 (GLP1R antagonist, 100 mg/kg). Data are represented as mean ± SEM. n= 7 – 9 biological replicates for each genotype. *p<0.05 between genotypes within time by Student’s t test.

### 3.6. Fasting Produces Greater Increases in Acute Food Intake in LPGC1a+/- Mice

Acute reduction of liver ATP levels by 2,5-anhydrous mannitol injection has been shown to stimulate food intake [41; 42; 53-55]. Further, intact vagal nerve function below the common hepatic branch was necessary to elicit the increased food intake following injection [41; 42]. However, the intraperitoneal delivery route of 2,5-anhydrous mannitol and the complete loss of efferent and afferent vagal activity below the common hepatic branch following vagotomy limit the interpretation of these findings. The reduced liver ATP concentrations following an overnight fast observed in the male LPGC1a+/- provide a unique opportunity to assess the role of hepatic energy status in acute food intake regulation. In a different set of mice, an 18 hr fast increased acute cumulative food intake (65%) in WT mice compared to their food intake during 2hr food withdrawal control experiments (**Figure 6A**). However, male LPGC1a+/- mice had ∼1.5-times greater acute food intake as compared to their control consumption. This increase in food intake was primarily due to increased meal size observed in the first 30 minutes following access to chow in both genotypes (**Figure 6B**). Though the WT mice increased the size of meals during this period greater than 3-fold, the male LPGC1a+/- mice increased meal size greater than 13.5-fold. Further, the LPGC1a+/- mice continued to have larger meal size (>2.5 fold) during the following 30-minute period, while the WT had returned to control values. This increase in food intake was not due to an increase in either the length of meals per period (**Figure 6C**) or the number of meals per period (**Figure 6D**). Further, LPGC1a+/- fasting-induced food intake was not as sensitive to peripheral satiation signal inhibition with CCK-8S (10 μg/kg) compared to WT (**Figure 6E & F**). While these findings are striking, it must be noted that no difference in control meal size or meal duration were observed in these experiments unlike the previous findings above (**Figure 1**).

**Figure 6.**
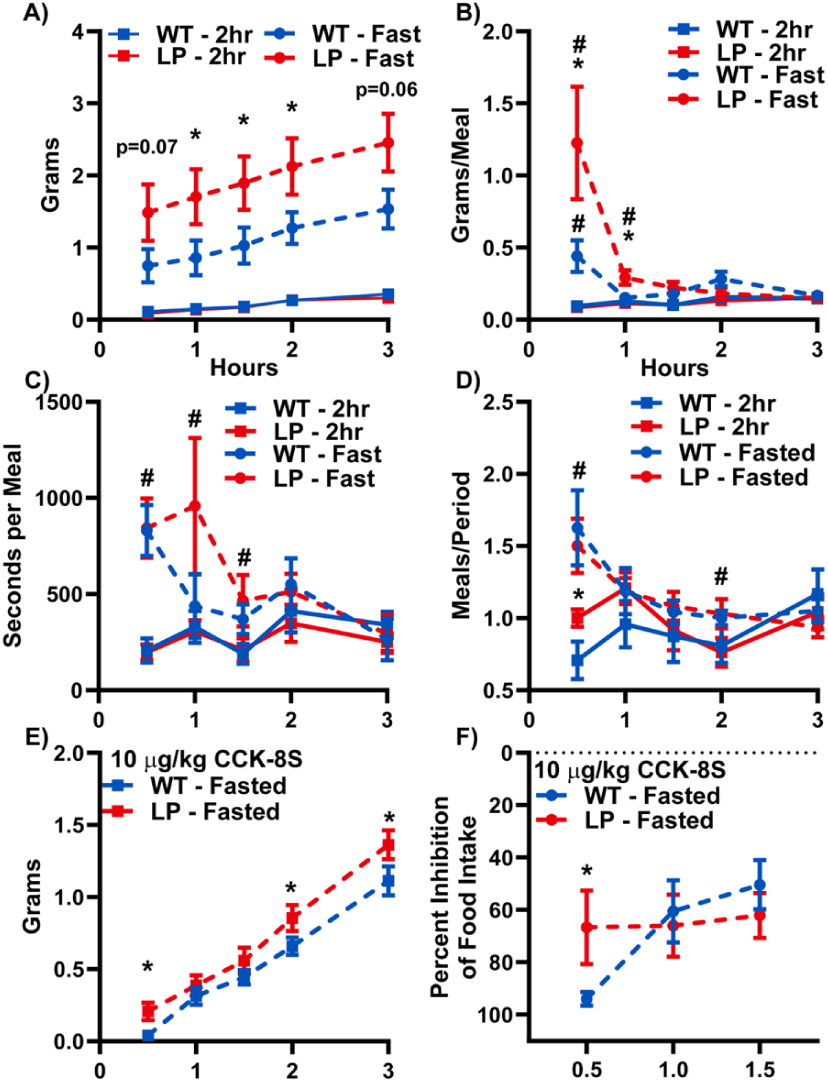
Fasting Produces Greater Increases in Acute Food Intake in LPGC1a+/- Mice. A) Cumulative food intake, B) grams/meal per 0.5 hr period, C) meals per period, and D) seconds per meal in WT and LPGC1a+/- mice fasted for 18 hr compared to 2hr food withdrawal. E) Cumulative food intake and F) percent inhibition of food intake in 18hr fasted mice receiving 10 mg/kg CCK-8S prior to access to food. Data are represented as mean ± SEM. n= 7 – 9 biological replicates for each genotype. *p<0.05 between genotypes within time and # p<0.05 between treatment within genotype by Student’s t test.

These data demonstrate a robust association of fasting-induced reductions in liver ATP and increases in acute food intake. However, the liver secretes numerous hepatokines that are regulated by fasting/re-feeding and/or regulate food intake [94]. We assessed the gene expression of 6 hepatokines (i.e., FGF21, GDF15, LEAP2, Angptl4, FetuA, and FetuB) thought to be involved in food intake regulation in fasted/re-fed male WT and LPGC1a+/- mice (**Figure S7**). Only FGF21 expression was lower in LPGC1a+/- mice during fasting (p=0.12). While, liver FGF21 expression has been observed to be increased in fasting, it’s role in food intake regulation has been refined to increased preference of protein and inhibition of carbohydrate consumption[94]. Previously, we have observed similar FGF21 serum levels in WT and LPGC1a+/- mice [95], however, these mice were not fasted overnight. Future experiments may investigate a potential role of reduced fasting liver FGF21 expression in acute food intake. Together, these findings demonstrate that the LPGC1a+/- mice have exaggerated acute food intake following an overnight fast, mainly through the consumption of larger meals.

### 3.7. Maladaptive Response of Hypothalamic POMC and AgRP Expression to Fasting in LPGC1a+/- Mice

Orexigenic AgRP/NPY and anorexigenic POMC neurons of the arcuate nucleus are central to the homeostatic regulation of food intake, with fasting serving as a powerful modulator of their expression. We measured the gene expression of these neuropeptides in the hypothalamus of fasted/re-fed male WT and LPGC1a+/- mice to determine whether the observed differences in fasted food intake were associated with differences in these homeostatic genes. Male WT mice were observed to have an expected decrease in POMC during fasting and subsequent increase following feeding (**Figure 7A**). Interestingly, POMC expression was lower in LPGC1a+/- mice under basal conditions and no change in expression was observed during either fasting or re-feeding. Importantly, AgRP expression was higher in LPGC1a+/- mice compared to WT during fasting (**Figure 7B**), while NPY expression was increased with fasting in WT but not LPGC1a+/- mice (**Figure 7C**). Based on these findings, we next assessed the total area of POMC and AgRP transcript expression in the arcuate nucleus by *in situ* hybridization in mice after a 4-hour food withdrawal. We observed a reduction in POMC (**Figure 7D**) and an increase in AgRP fluorescent area (**Figure 7E**) in the male LPGC1a+/- mice. However, the increase in fasting-induced acute food intake in Figure 6 is consistent with increased expression of AgRP in the hypothalamus of male LPGC1a+/- mice following fasting. Together, fasting reduces liver ATP, increases hypothalamic orexigenic gene expression, and elevates acute food intake in LPGC1a+/- mice.

**Figure 7.**
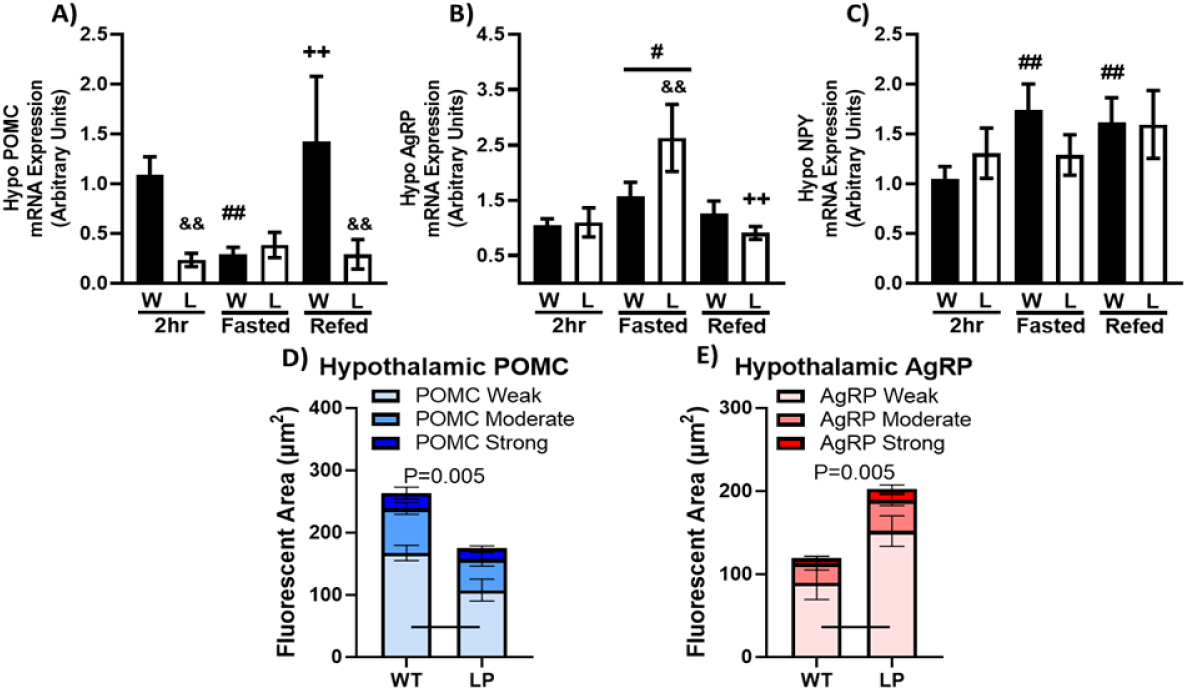
Maladaptive Response of Hypothalamic POMC and AgRP Expression to Fasting in LPGC1a+/- Mice. Gene expression of A) POMC, B) AgRP, and C) NPY in the hypothalamus of 2hr food withdrawn, fasted, and 4 hr refed mice. *In situ* hybridization of D) POMC and E) AgRP in the arcuate nucleus of the hypothalamus of 2hr food withdrawn wildtype and LPGC1a+/- mice. Data are represented as mean ± SEM. n= 7 – 9 biological replicates for each genotype. # main effect fasting compared to 2hr by two-way ANOVA (B). ## fasting versus 2 hr within genotype, && LPGC1a+/- versus wildtype within group, and ++ refed versus fasting within genotype pairwise comparisons were performed using Fishers LSD (A, B, & C). D) and E) genotyped compared by Student’s t test.

## 4. Discussion

In the 1960s, Russek first suggested that liver metabolism could serve as a sensor in the peripheral regulation of food intake [39; 96-98]. Subsequently, Langhans and Friedman independently reported vagal afferent communication of liver MEM to higher neural structures as a mediator of food intake. These studies chemically blocked fatty acid oxidation (FAO) [44-46; 50-52] and/or lowered ATP levels [41; 42; 53-56] which increased acute food intake, altered meal patterns [47; 99], and was hypothesized to decrease vagal afferent activation in rodents [42; 43; 100-102]. Furthermore, common hepatic branch vagotomy or chemical deafferentation prevented the effects of reduced ATP or FAO [40-43; 103; 104]. However, it was later proposed that sensing of small intestine FAO, not hepatic, was involved in peripheral food intake regulation [105; 106]. This premise, combined with prior research showing a lack of vagal afferent innervation of the liver parenchyma [107] and the reduced rigor of intraperitoneal chemical inhibition and vagotomy experiments, reduced interest in the study of liver MEM as a mediator of food intake. However, numerous recent findings coupled with major technological advancements have revived the hypothesis that liver MEM can modulate vagal afferent sensory neuron signaling impacting energy homeostasis [62; 108-110] and metabolic disease development and progression [85; 86; 111-114]. Herein, we report impaired food intake inhibition following and oral nutrient pre-load and greater fasting-induced food intake in a mouse model with decreased hepatocyte MEM (e.g., oxidative metabolism [62], respiratory function, adaptability to changes in energy demand, and ATP homeostasis). These food intake phenotypes were characterized by greater within meal food intake and meal length, suggesting impaired GI-derived satiation signaling. In support of compromised satiation signaling, peripheral antagonism of vagal afferent satiation signal receptors failed to increase relative food intake following oral nutrient pre-load. Also, fasting-induced food intake was not as strongly inhibited by peripheral delivery of a satiation signal .

A large body of work suggests that altered liver mitochondrial oxidative metabolism and ATP homeostasis is associated with obesity. In human subjects with obesity, liver ATP concentration is inversely proportional to BMI [115–117], waist-to-hip ratio [118; 119], and hepatic lipid content [115]. Importantly, obesity drives compensatory responses in hepatic oxidative metabolism [120], including increased TCA flux [121–123]. However, these adaptations do not protect liver ATP homeostasis in obese individuals as observed by reduced basal or fasting hepatic ATP concentration [116; 119; 124], ATP recovery rate [116; 125-127], and poor coupling of mitochondrial oxygen consumption to ATP synthesis in isolated mitochondria (as respiratory control ratio) [128]. This reduction in tissue ATP homeostasis is due in part to decreased electron transport system (ETS) complex activities [128; 129], complete FAO to CO_2_ [130], mitochondrial turnover [130], and mitochondrial entry of long-chain fatty acids in subjects with obesity [120]. Similarly, DIO rodents demonstrate reduced expression [131–137] and activity [132-135; 137; 138] of ETS complexes, and, impaired capacity for maintaining hepatic ATP levels [132-135; 139]. In humans and rodents, these impairments in hepatic MEM are associated with decreased expression of the mitochondrial co-transcriptional regulator, peroxisomal proliferator-activated receptor γ co-activator 1α (PGC1a) [124; 128; 132-134; 136; 139-146], as well as, its transcriptional targets involved in mitochondrial transcription pathways: mitochondrial transcription factor-a (TFAM) [137; 139; 141; 142] and nuclear respiratory factor-1 [139; 141]. This association of obesity, impaired liver MEM, and the reduced satiation observed in obesity implicates liver ATP homeostasis as a potential regulator of the interoceptive processes leading to within-meal food intake inhibition. Importantly, the LPGC1a+/-model recapitulates the impaired MEM (**Figure 1**), particularly the loss of ATP homeostasis, without the potential confounding effects of diet-induced obesity. These findings, together with the observed impaired satiation in obesity [19–35], implicate liver mitochondrial MEM in the regulation of food intake in overweight and obese pre-clinical models and humans.

Previously, we have observed increased acute high-fat diet food intake and greater weight gain in animals with chronically reduced liver MEM [58–60]. Alternatively, elevated hepatic FAO via CPT1a overexpression decreased food intake during chronic high-fat diet feeding [147]. Further, increased liver glucose flux resulted in decreased weight gain, reduced vagal-dependent reductions in food intake, and increased orexigenic gene expression in the hypothalamus [108–110]. As described above, early data implicated liver MEM in the regulation of food intake via modulation of vagal afferent neuron signaling through chemical inhibition of liver fatty acid oxidation and/or ATP depletion [40; 42-45; 51; 52; 54; 55; 57; 100-102; 104]. Recently, we observed that the reduced liver MEM of male LPGC1a+/- mice was associated with increased short-term high-fat diet food intake, larger meal size, and greater weight gain compared to WT littermates [62]. Importantly, no differences were observed in food intake or weight gain between LPGC1a+/- and WT mice on low-fat diet. While the increased high-fat diet food intake and meal size suggested impaired food intake regulation, the pathophysiology was unknown. In the current study, we observed LPGC1a+/- mice have impaired underlying differences in meal size, duration, and number without differences in food intake during normal chow feeding (**Figure 2**). Further, reduced liver MEM in LPGC1a+/- mice was associated with decreased dose-dependent food intake inhibition by oral pre-loads (**Figure 3**), lack of increase in food intake following antagonism of peripheral GI-derived satiation signal receptors (**Figure 5**), and increased fasting-induced food intake (**Figure 6**). The reduced mitochondrial fatty acid oxidation [62], respiration of FFA (**Figure 1A**), and impaired ATP homeostasis (**Figure 1B**) of the male LPGC1a+/- mouse replicated the reduced liver energy metabolism of the earlier studies, while more specifically targeting the liver. To our knowledge this is the first data showing the potential role of liver MEM on within meal feeding behavior and food intake inhibition.

Our findings suggest a role for liver MEM in the interoceptive control of meal termination by GI-derived satiation signals. Appropriate meal termination requires higher order brain structures to integrate neuroendocrine signals initiated by gastric distension and gut-derived neuropeptides/neurotransmitter activation of vagal afferent neurons (reviewed in [148–151]). The modulation of these neural signals by peripheral energy metabolism could serve as a mechanism to communicate peripheral tissue energy state to the brain. Recently, Geisler et al. proposed that reduced hepatic ATP lowers Na/K-ATPase activity, resulting in hepatocyte membrane depolarization, increased GABA secretion, and inhibition of vagal afferent firing [85; 86]. As described above, liver ATP levels are reduced in human subjects and rodent models with obesity. Additionally, liver Na/K-ATPase activity is reduced in diet-induced obese rodents [152], and IP delivery of ouabain (Na/K-ATPase inhibitor) increases food intake [153]. Further, chemical inhibition of fatty acid oxidation [154], Na/K-ATPase activity [155–158], ATP depletion [157], and electrogenic nutrient uptake [159] depolarizes primary hepatocytes. Classically, the liver is considered a primary site of peripheral GABA degradation through GABA-transaminase and the GABA shunt, as such, hepatocytes express four GABA transporters which are involved in sodium-dependent GABA uptake [160]. However, the equilibrium constant of GABA-transaminase favors GABA synthesis [161]. Hepatocyte GABA synthesis through GABA-T would be favored by increased generation of succinate semialdehyde through the low liver NAD+/NADH observed in fasted male LPGC1a+/- mice (**Figure 1D**) and obese models [85; 132; 133]. As such, GABA secretion is increased in obese mouse liver sections and following inhibition of Na/K-ATPase [85]. Finally, using hepatocyte-specific expression of a ligand-gated sodium channel, the authors demonstrated real-time reductions in vagal afferent firing rate and hepatocyte membrane depolarization during ligand application [85]. The maintenance of hepatocyte membrane potential by ATP-dependent Na/K-ATPase activity represents ∼35% of hepatocyte oxygen consumption [89]. As such, any reduction in liver ATP homeostasis could lead to hepatocyte membrane depolarization and reduced vagal afferent communication of peripheral satiation signals. Our observations of increased acute meal size (**Figure 6B**) and impaired CCK inhibition of food intake (**Figure 6F**) in fasted LPGC1a+/- mice with reduced liver ATP supports this speculation. Future studies will use hepatocyte-specific expression of newly developed ultrapotent ligand-gated channels to target experiments on the role of hepatocyte membrane polarization state in meal termination by GI-derived satiation signals.

A minor number of limitations require consideration in the interpretation and translation of our findings. One, in our previous work we did not observe differences in diet-induced weight gain or high-fat diet food intake in female wildtype and LPGC1a+/- mice [62]. While this is why we have focused on males to this point, we are unable to discuss the role of sex on liver MEM and food intake regulation. Future work will include female groups to assess if liver MEM is involved in the previously observed phenotypes. Two, hepatocyte PGC1a co-regulates other transcriptional pathways beyond mitochondrial hormesis, particularly, gluconeogenesis, antioxidant, and anti-inflammatory pathways [82]. As such, it is possible that changes in these transcriptional pathways in hepatocytes are influencing the observed food intake regulation phenotypes. Although, LPGC1a+/- mice had normal nutritional regulation of PEPCK expression, suggesting gluconeogenic activity was not altered. Future work will focus on more specific modeling of reductions in hepatocyte mitochondrial ATP homeostasis. Three, limitations of the rigor of the modeling system must be considered. Limited data suggests that albumin is expressed in non-hepatic tissues [162; 163], which would impact the tissue-specificity of the PGC1a heterozygosity. Further, expression of albumin-*cre* recombinase begins approximately ∼half way through gestation (∼10.5 of 20 day gestation [164], potentially resulting in developmental changes that impact adult metabolic physiology. This is supported by the findings that offspring of high-fat fed dams have decreased liver PGC1a expression [165; 166], and increased AgRP/NPY and decreased POMC hypothalamic expression [167]. Future experiments will be performed in adult onset models using hepatocyte-specific delivery of cre recombinase via AAV.

## Conclusions

In summary, we demonstrate that decreased liver MEM alters feeding microstructure and food intake inhibition in response to nutrients. In combination with others[85; 86], these data support a novel interoceptive mechanism by which liver energy metabolism can modulate acute food intake through reduced vagal afferent activity due to increased hepatocyte GABA secretion secondary to impaired maintance of membrane potential. Experiments are ongoing to further investigate the capacity of hepatic mitochondrial energy metabolism and ATP homeostasis to modulate meal termination in male and female mice through 1) increased rigor and specificity of hepatic mitochondrial models, 2) assessment of hepatocyte membrane potential and GABA under conditions of reduced hepatocyte MEM, and 3) determination of food intake regulation during acute chemogenetic hepatocyte membrane potential manipulation. Future work will include confirming the necessity of vagal afferents in the regulation of food intake through transmission of reduced hepatocyte MEM, and identifying the vagal afferent and higher order neural populations involved.

## Resource Availability

Requests for further information and resources should be directed to and will be fulfilled by the lead contact, E. Matthew Morris.

## Author Contributions

Author contributions: EMM, CEG, MRH, Conceptualization; EMM, Data curation; MEP, JCP, MAN, ABD, LLC, HH, CEG, EMM, Formal analysis; MEP, JCP, MAN, ABD, LLC, HH, CEG, MRH, EMM, Writing – original draft; MEP, JCP, MAN, ABD, LLC, HH, CEG, MRH, EMM, Writing editing and revising.

## Acknowledgements

The authors wish to thank David Wasserman for years of scientific mentoring, career advice, and comedy relief. You will be missed.

## Supplemental Information

Figures S1-S7

