## Supplemental Figures for "Reduced Liver Mitochondrial Energy Metabolism Impairs Food Intake Regulation Following Gastric Preloads and Fasting"

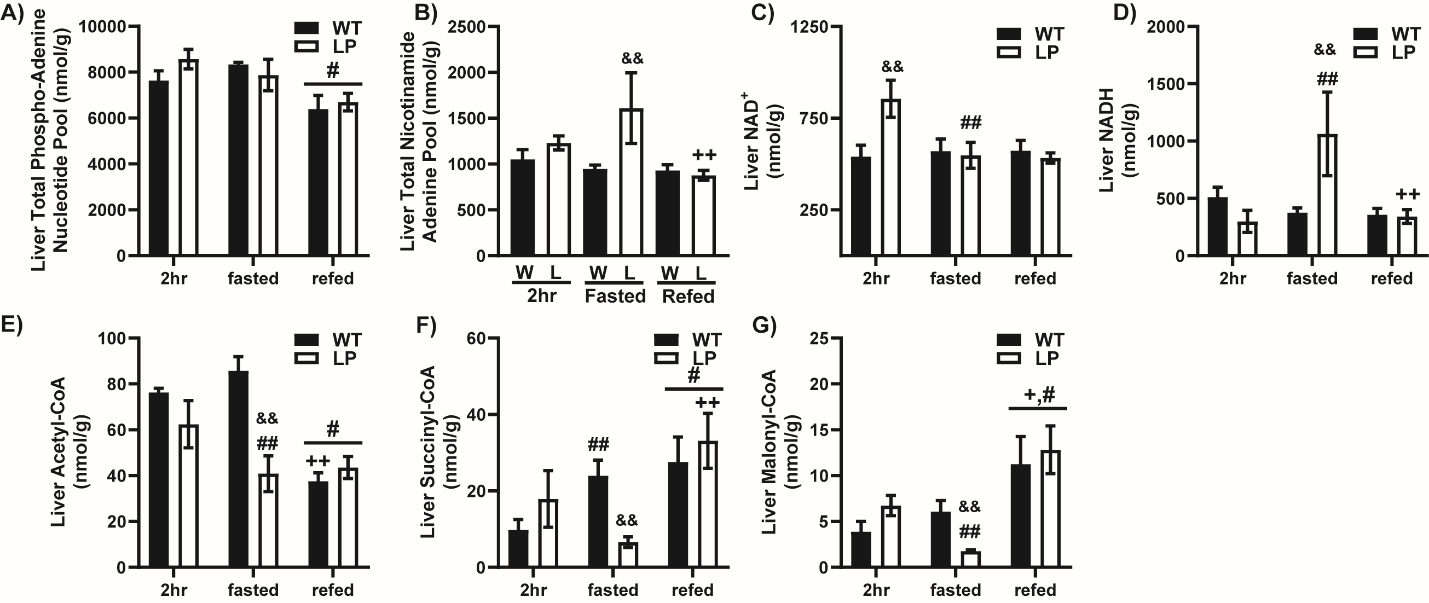
**Figure S1**

**Figure S1. Liver Adenine Nucleotides during Fasting/Refeeding.** Livers from male WT and LPGC1a+/- mice following 2 hr food withdrawal, 18 hr fast, or 4 hr refeeding were used to quantify A) total phosphor-adenine nucleotide pool, B) total nicotinamide adenine pool, C) NAD+, D) NADH, E) acetyl-CoA, F) succinyl-CoA, and G) malonyl-CoA. Data are represented as mean ± SEM. n= 3 – 6 biological replicates for each genotype. # main effect fasting or refeeding compared to 2hr, & main effect of genotype, + main effect of refed vs fasted by two-way ANOVA. *p<0.05 between genotypes by Student’s t-test (A). # main effect vs 2hr, + main effect refed vs fasted, & main effect LPGC1a+/- vs WT. ## fasting versus 2 hr within genotype, && LPGC1a+/- versus wildtype within group, and ++ refed versus fasting within genotype pairwise comparisons were performed using Fishers LSD.

**Figure S2**

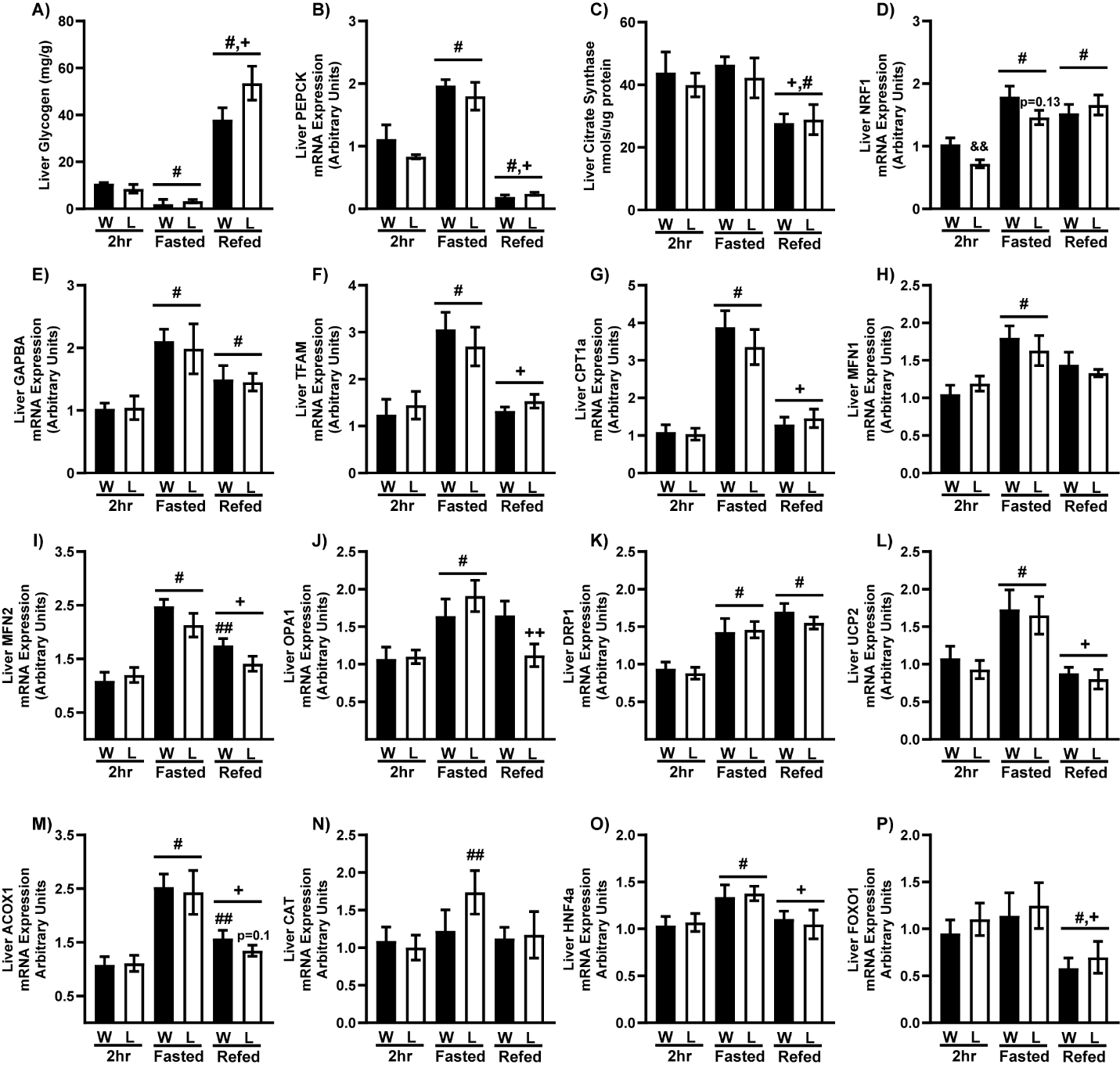

**Figure S2. Liver Mitochondrial Gene Expression during Fasting/Refeeding.** The impact of fasting/refeeding on liver A) glycogen concentration, B) PEPCK mRNA, and C) citrate synthase activity was determined in male WT and LPGC1a+/- after 2 hr food withdrawal, 18 hr fast, or 4 hr refeeding. Livers from male WT and LPGC1a+/- mice following 2 hr food withdrawal, 18 hr fast, or 4 hr refeeding were used to determine gene expression of D) NRF1, E) GAPBA, F) TFAM, G) CPT1a, H) MFN1, I) MFN2, J) OPA1, K) DRP1, L) ESRR alpha, M) ACOX1, N) CAT, O) HNF4a, and P) FOXO1. Data are represented as mean ± SEM. n= 8 – 10 biological replicates for each genotype. # main effect fasting or refeeding compared to 2hr, & main effect of genotype, + main effect of refed vs fasted by two-way ANOVA. *p<0.05 between genotypes by Student’s t-test (A). # main effect vs 2hr, + main effect refed vs fasted, & main effect LPGC1a+/- vs WT. ## fasting versus 2 hr within genotype, && LPGC1a+/- versus wildtype within group, and ++ refed versus fasting within genotype pairwise comparisons were performed using Fishers LSD.

**Figure S3**

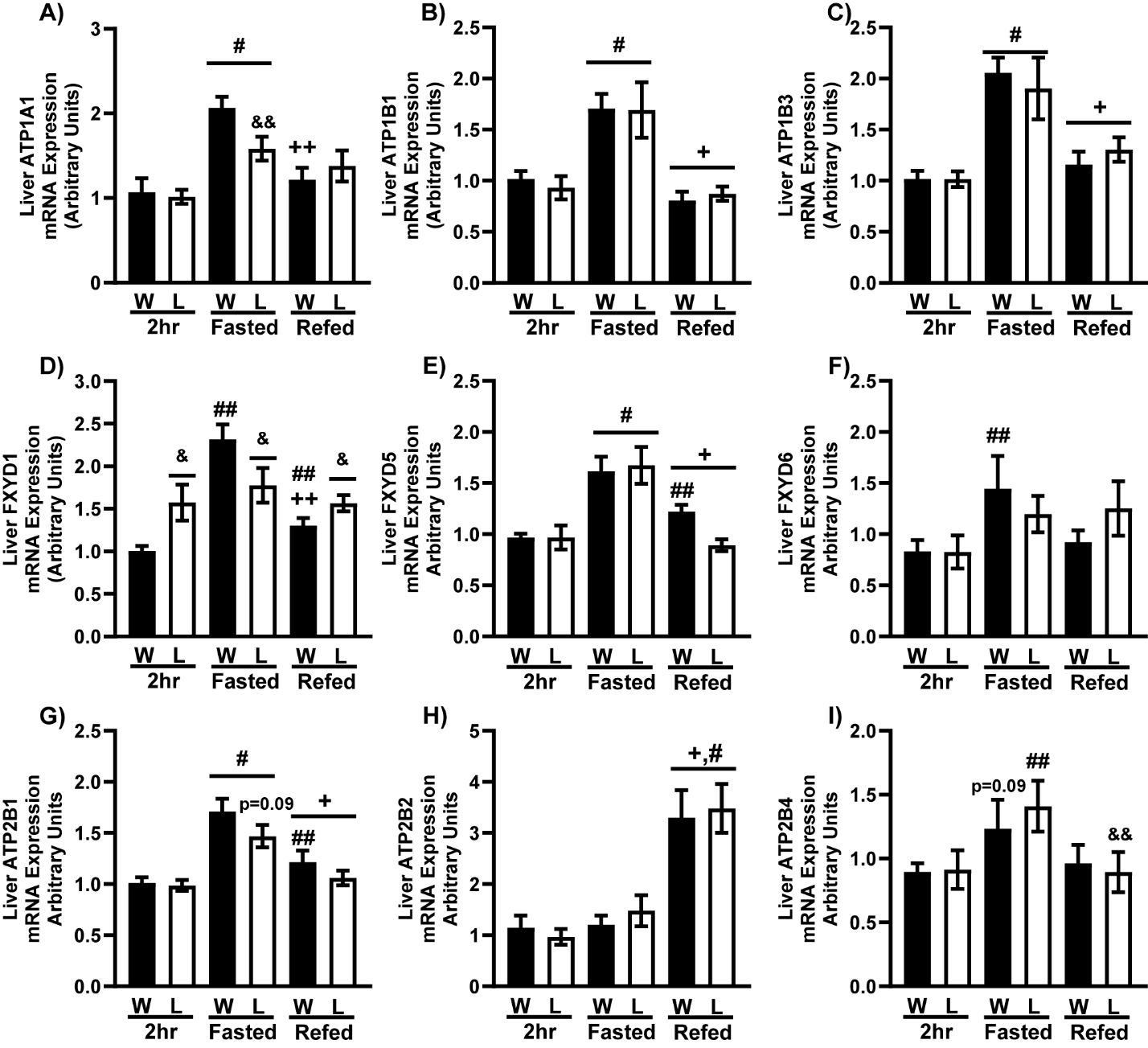

**Figure S3. Liver Na/K-ATPase Gene Expression during Fasting/Refeeding.** Livers from male WT and LPGC1a+/- mice following 2 hr food withdrawal, 18 hr fast, or 4 hr refeeding were used to determine gene expression of A) ATP1a1, B) ATP1B1, C) ATP1B3, D) FXYD1, E) FXYD5, F) FXYD6, G) ATP2B1, H) ATP2B2, and I) ATP2B4.. Data are represented as mean ± SEM. n= 8 – 10 biological replicates for each genotype. # main effect fasting or refeeding compared to 2hr, & main effect of genotype, + main effect of refed vs fasted by two-way ANOVA. *p<0.05 between genotypes by Student’s t-test (A). # main effect vs 2hr, + main effect refed vs fasted, & main effect LPGC1a+/- vs WT. ## fasting versus 2 hr within genotype, && LPGC1a+/- versus wildtype within group, and ++ refed versus fasting within genotype pairwise comparisons were performed using Fishers LSD.

**Figure S4**

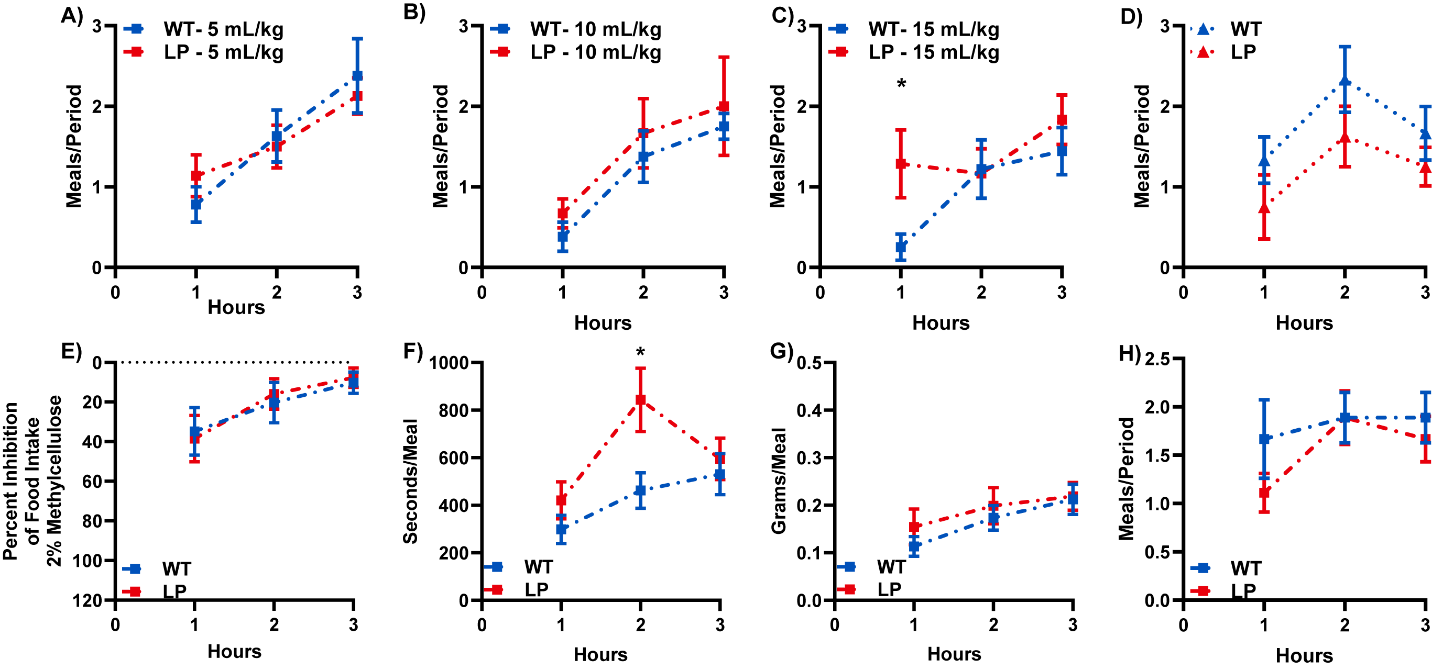

**Figure S4. Meals per Period following Mixed Nutrient Oral Gavage Pre-Loads.** Meals per period following A) 5 mL/kg, B) 10 mL/kg, C) 15 mL mL/kg, and D) 10 ml/kg of 1% methylcellulose oral gavage pre-load in male WT and LPGC1a+/- mice. E) Percent food intake inhibition, F) seconds per meal, G) grams per meal, and H) meals per period in male WT and LPGC1a+/- mice following 10 mL/kg oral gavage pre-load of 2% methylcellulose. Data are represented as mean ± SEM. n= 7 – 9 biological replicates for each genotype. *p<0.05 between genotypes within time and # p<0.05 between treatment within genotype by Student’s t test.

**Figure S5**

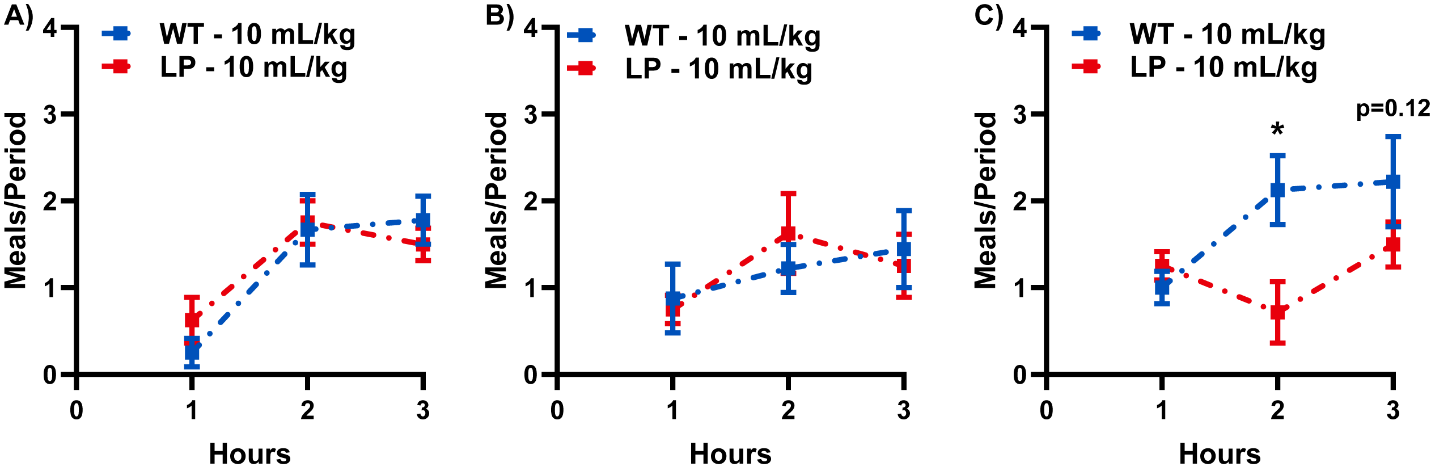

**Figure S5. Meals per Period following Individual Nutrient Oral Gavage Pre-Loads.** Meals per period following 10 mL/kg oral gavage pre-load of A) Proteinex, B) Intralipid, and C) 40% glucose in male WT and LPGC1a+/- mice. Data are represented as mean ± SEM. n= 7 – 9 biological replicates for each genotype. *p<0.05 between genotypes within time and # p<0.05 between treatment within genotype by Student’s t test.

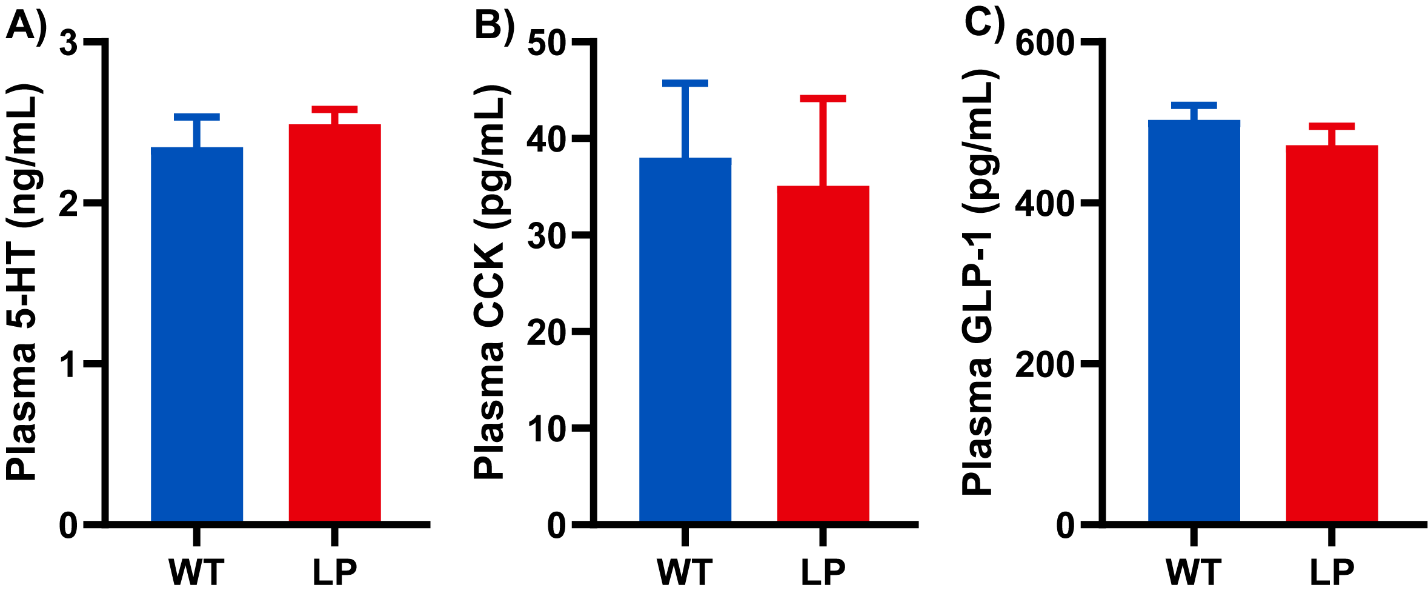
**Figure S6**

**Figure S6. Meals per Period following Individual Nutrient Oral Gavage Pre-Loads.** Plasma concentration of A) 5-HT, B) CCK, and C) GLP-1 in male WT and LPGC1a+/- mice 60 minutes following a 10 mL/kg Ensure oral gavage. Data are represented as mean ± SEM. n= 7 – 9 biological replicates for each genotype. *p<0.05 between genotypes within time and # p<0.05 between treatment within genotype by Student’s t test.

**Figure S7**

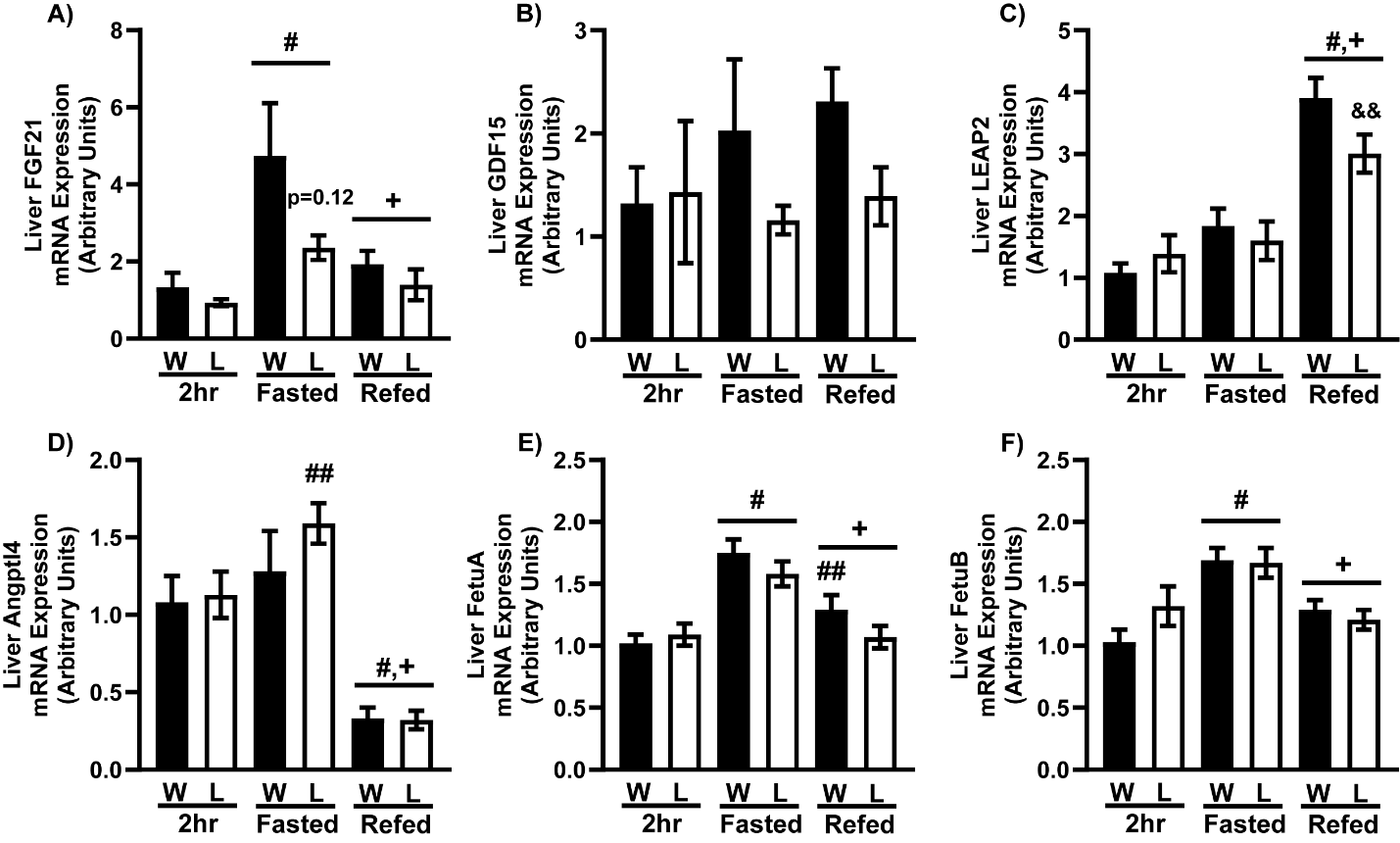

**Figure S7. Liver Hepatokine Gene Expression during Fasting/Refeeding.** The impact of fasting/refeeding on liver mRNA expression of A) FGF21, B) GDF15, C) LEAP2, D) Angptl4, E) FetuA, and F) FetuB was determined in male WT and LPGC1a+/- after 2 hr food withdrawal, 18 hr fast, or 4 hr refeeding. Data are represented as mean ± SEM. n= 8 – 10 biological replicates for each genotype. # main effect fasting or refeeding compared to 2hr, & main effect of genotype, + main effect of refed vs fasted by two-way ANOVA. *p<0.05 between genotypes by Student’s t-test (A). # main effect vs 2hr, + main effect refed vs fasted, & main effect LPGC1a+/- vs WT. ## fasting versus 2 hr within genotype, && LPGC1a+/- versus wildtype within group, and ++ refed versus fasting within genotype pairwise comparisons were performed using Fishers LSD.

**Figure S8**

**Figure S8. Representative Fluorescent In Situ Hybridization Images.** Representative images of the fluorescent in situ hybridization of POMC (green) and AgRP (red) in the arcuate nucleus of male WT and LPGC1a+/- mice: A) WT rostral arcuate nucleus, B) LPGC1a+/- rostral arcuate nucleus, C) WT caudal arcuate nucleus, and D) LPGC1a+/- caudal arcuate nucleus.

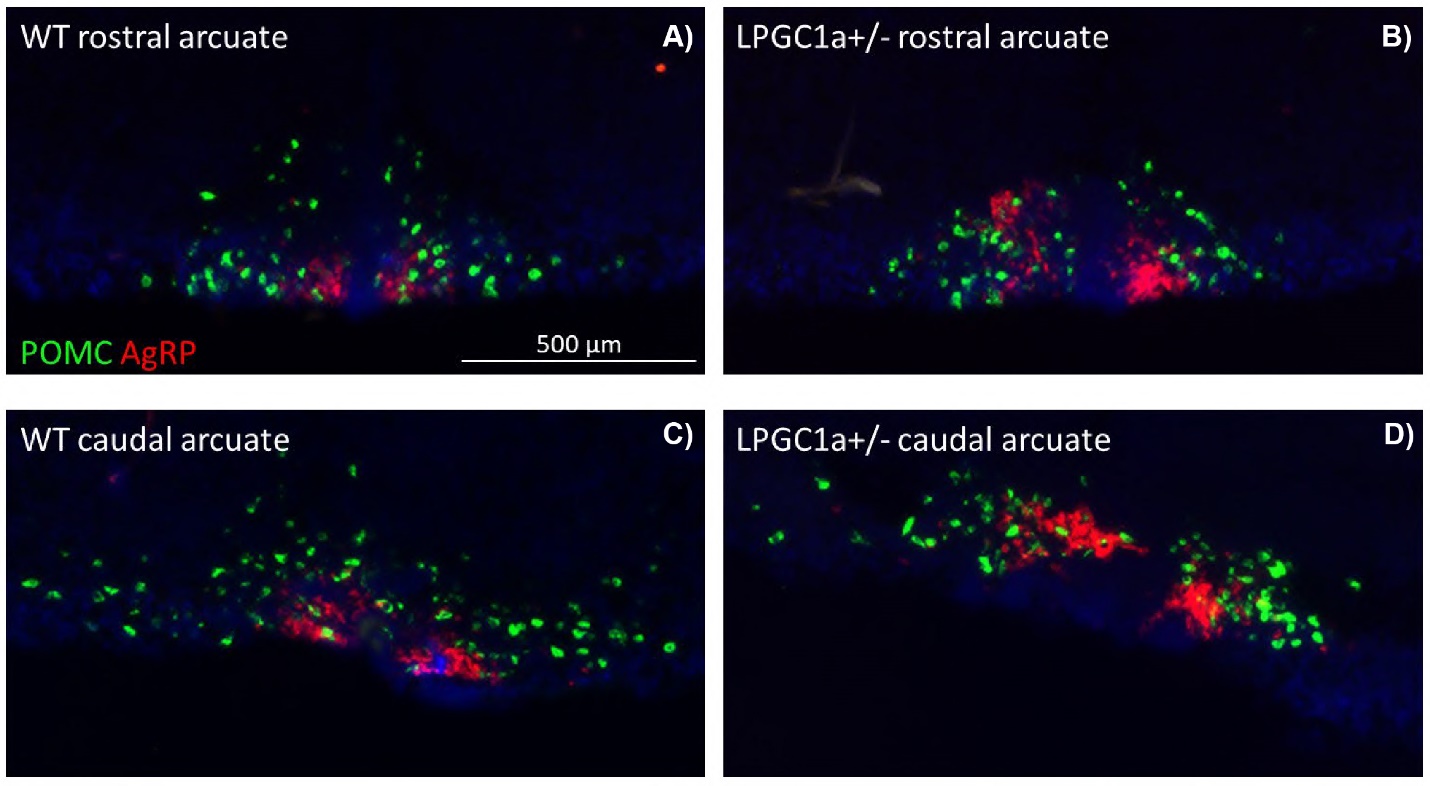
